# A single-component luminescent biosensor for the SARS-CoV-2 spike protein

**DOI:** 10.1101/2022.06.15.496006

**Authors:** Matthew Ravalin, Heegwang Roh, Rahul Suryawanshi, G. Renuka Kumar, John Pak, Melanie Ott, Alice Y. Ting

## Abstract

Many existing protein detection strategies depend on highly functionalized antibody reagents. A simpler and easier to produce class of detection reagent is highly desirable. We designed a single-component, recombinant, luminescent biosensor that can be expressed in laboratory strains of *E. coli* and *S. cerevisiae*. This biosensor is deployed in multiple homogenous and immobilized assay formats to detect recombinant SARS-CoV-2 spike antigen and cultured virus. The chemiluminescent signal generated facilitates detection by an un-augmented cell phone camera. <u>B</u>inding <u>A</u>ctivated <u>T</u>andem split-enzyme (BAT) biosensors may serve as a useful template for diagnostics and reagents that detect SARS-CoV-2 antigens and other proteins of interest.

## Introduction

The detection and quantification of analytes in complex biological samples is essential for both discovery and translational biomedical research. Small-molecule and protein analyte detection are dominated by antibody-based technologies. Such technologies have three major limitations: 1) antibodies are generally produced in mammalian expression systems that require significant infrastructure, 2) nearly all antibody-based assays require chemical modification to immobilize the antibodies or conjugate detection reagents, and 3) these platforms require at least two and up to four unique antibodies to function. To address these limitations, we and others have developed alternative biosensor platforms that can be more easily produced and work in both homogenous and immobilized assay formats.^1^

Split luciferase enzymes are attractive for biosensor design because they are highly sensitive and, in contrast to fluorescent reporters that require an excitation source, need only a camera for detection. Several two-component luciferase-based biosensors have been reported, in which analyte binding drives the reconstitution of the split enzyme (**Figure S1a, b**).^2–8^ A challenge with such sensors is the requirement for two separate binding modules that interact with independent epitopes on the target analyte. Recently, the LOCKR platform used a single binding module grafted into a caging structure that allows for enzyme reconstitution triggered by analyte binding. (**Figure S1c**).^9–11^ In this system, the binding module serves as the “lock” for a caging structure that masks half of split luciferase. The analyte serves as a “key” that opens the cage and allows the association of the other half of the luciferase which is added as a second component.

Single-component luminescent biosensors are potentially simpler to produce and easier to use. Existing designs have relied on naturally-occurring binding modules that undergo large conformational changes upon binding (**Figure S1d**)^12–14^ or tethered decoy analytes that stabilize an auto-inhibited state of the sensor (**Figure S1e**).^15^ These approaches are difficult to generalize and limit the scope of analytes that can be detected.

Here we introduce a single-component, NanoLuc-based, <u>B</u>inding <u>A</u>ctivated <u>T</u>andem split enzyme (BAT) biosensor. It does not rely on a large conformational change in the binding module or competition with a tethered decoy as with other single component platforms. We use this luminescent BAT biosensor for the detection of the SARS-CoV-2 spike protein in multiple assay formats.

## Results

### Design and optimization of SARS-CoV-2 biosensor “S-BAT”

In our design, a small, rigid binding module is fused to split luciferase halves in such a fashion as to disfavor their reconstitution, due to unfavorable geometry and low effective concentration. Antigen binding introduces steric clashes with the luciferase halves that re-orient them and increase their effective concentration, driving reconstitution (**Figure 1a, Figure S1f**). To demonstrate the BAT concept, we selected the computationally designed miniprotein LCB1 as our binder. LCB1 binds to the receptor binding domain (RBD) of SARS-CoV-2 spike protein with a K_d_ of 10^-10^ M.^16^ In addition to having high thermal stability, rigidity, and no disulfide bonds to complicate purification, the N- and C-termini in the three-helix bundle scaffold project in opposite directions (∼25 Å apart, based on measurements from PDB ID 7JZU), providing a rigid scaffold for minimizing the effective concentration of the fused split luciferase halves.

**Figure 1.**
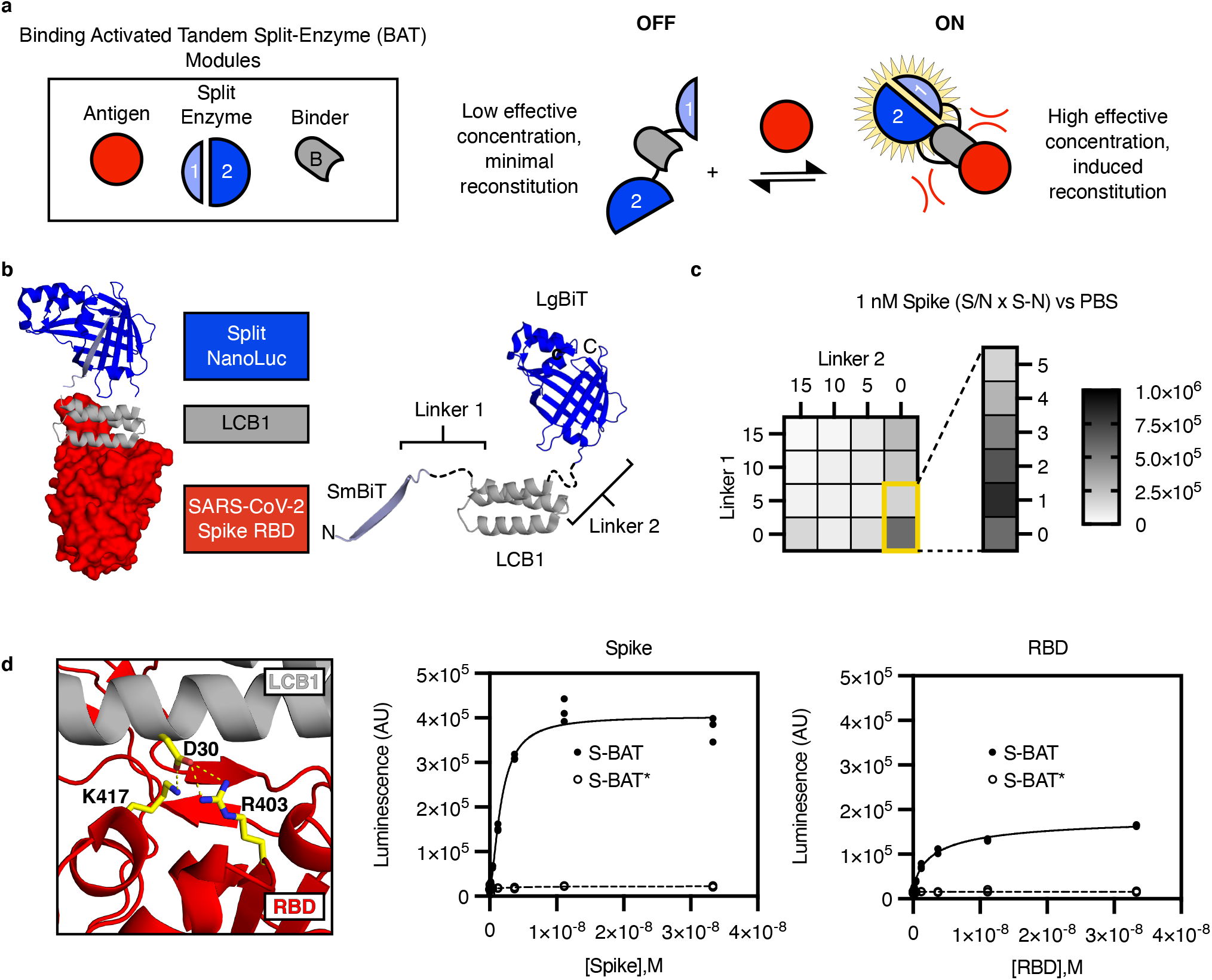
Design of a binding activated tandem split enzyme (BAT) biosensor for the SARS-CoV-2 Spike protein. a) BATs consist of a binding module (B, grey) for antigen (red) and a split enzyme (1 & 2, Blue). The split enzyme is fused in tandem to the N and C terminus of the binding module. A low effective concentration of 1 and 2 in the “OFF” state limit background activity. Binding-induced steric clashes between the antigen and the tethered split enzyme components increase the effective concentration of 1 and 2, driving reconstitution in the “ON” state. b) A model of an activated BAT (left) composed of SmBiT (light blue, cartoon), the mini protein LCB1 (grey, cartoon), and LgBiT (blue, cartoon) bound to the receptor binding domain (RBD, red, surface) of SARS-CoV-2 spike protein. The lengths of Linker1 and Linker 2 (left, black, dashed) are likely to control the effective concentration of SmBiT and LgBiT in the absence of the binder. c) A heat map of BAT performance as a function of linker length. Performance (Signal to Noise (S/N) multiplied by the magnitude of signal change (S-N)) for each linker composition is plotted based on detection of 1 nM recombinant Spike in phosphate buffered saline (PBS). The best linker combination was a single amino acid (Gly) for Linker 1 and 0 length linker for Linker 2. This molecule is named S-BAT. d) Asp 30 in LCB1 (grey, cartoon) makes critical contacts with Arg 403 and Lys 417 in the RBD of SARS-CoV-2 spike protein (red, cartoon). Mutation of Asp 30 to Ala yields S-BAT* which is not activated by recombinant Spike (middle panel) or RBD (right panel). Luminescence is plotted in arbitrary units (AU) versus antigen concentration (molar, M) for individual replicate samples (n=3) in a representative experiment.

We selected the split luciferase NanoBiT for BATs. NanoBiT is based on the bright, blue-emitting, ATP-independent, cysteine-free luciferase NanoLuc, and its SmBiT and LgBiT components have relatively low affinity for one another (K_d_∼190 μM).^17^ In a previous study, the tandem fusion FKBP12-SmBiT-Linker-LgBiT-FRB showed minimal background reconstitution in the absence of rapamycin.^18^ We performed a linker screen, varying the glycine-serine rich sequence between LCB1 and NanoBiT fragments between 0 and 15 amino acids. (**Figure 1b**) The initial library consisted of sixteen constructs (LCB1^Linker1 length, Linker2 length^), which were expressed in *E. coli* fused to a TEV protease-cleavable N-terminal His_6_ tag. TEV cleaved constructs were screened in a NanoGlo assay (Promega) for their ability to detect 1 nM (146.7 ng/mL) recombinant Spike derived from the WA1 strain of SARS-CoV-2 in a homogenous microplate assay. The data from this screen clearly indicated that the absolute change in signal (S-N) and the signal to noise ratio (S/N) increased as linker length *decreased*, with LCB1^0-0^ yielding the best composite performance (S/N x (S-N)). Based on apparent instability of LCB1^0-0^ and the dramatic increase in S/N between 5 and 0 linker lengths, we further optimized Linker 1 by varying the length from 1-5 with single amino acid resolution (**Figure 1c, Figure S2a, b**). These efforts yielded LCB1^1-0^ as our best SARS-CoV-2 Spike protein sensor (“S-BAT”).

We generated a point mutant in the S-BAT binding module at Asp30 (“S-BAT*”), designed to ablate salt bridges formed with Lys417 and Arg403 in the Spike receptor binding domain (RBD). S-BAT* exhibited low background activity consistent with its parent molecule but showed minimal activation upon incubation with either recombinant Spike or RBD, even at saturating concentrations (**Figure 1d**). We found that both trimeric Spike and monomeric RBD could activate S-BAT, with different absolute change in signal. This suggests that *cis* activation is likely the predominant source of signal in the assay, but we cannot rule out the contribution of a *trans* activating mechanism. In the *trans* mechanism, simultaneous binding to multiple protomers in a single Spike might increase the effective concentration, driving activation. However, the absence of a hook effect at super-stoichiometric concentrations of Spike binding sites to sensor copies supports a predominantly *cis* activation mechanism. The sufficiency of a monovalent interaction to activate these sensors makes BATs a promising scaffold for protein detection.

### S-BAT is functional in multiple assay formats

While homogeneous assays are convenient in a laboratory setting, immobilized assay formats can offer versatility in a variety of use settings. Previous work has shown that BRET-based biosensors can detect analytes of interest following adsorption onto cellulose paper, relying on a ratiometric output of luminescence and fluorescence for detection of analyte.^19^ We adsorbed our S-BAT sensor onto paper and used the change in luminescence intensity as our signal. With this approach, we were able to specifically detect low nanomolar concentrations of antigen using S-BAT, with S-BAT* serving as a negative control (**Figure 2a**). Adsorption-based immobilization is advantageous in that it requires no chemical modification to the protein reagent. However, chemical conjugation with biotin and subsequent immobilization on streptavidin allows for the concentration of the biosensor on the surface of a heterogenous assay matrix and can minimize loss of signal intensity inherent in the use of an opaque porous matrix like cellulose paper. Due to the absence of disulfide bonds in our constructs (in contrast to antibodies), we could introduce single cysteine residues throughout the protein as a handle for site specific bioconjugation. We generated five constructs containing a single cysteine residue and conjugated Biotin-PEG_n_-Maleimide (where n=2 or 11) to each.

**Figure 2.**
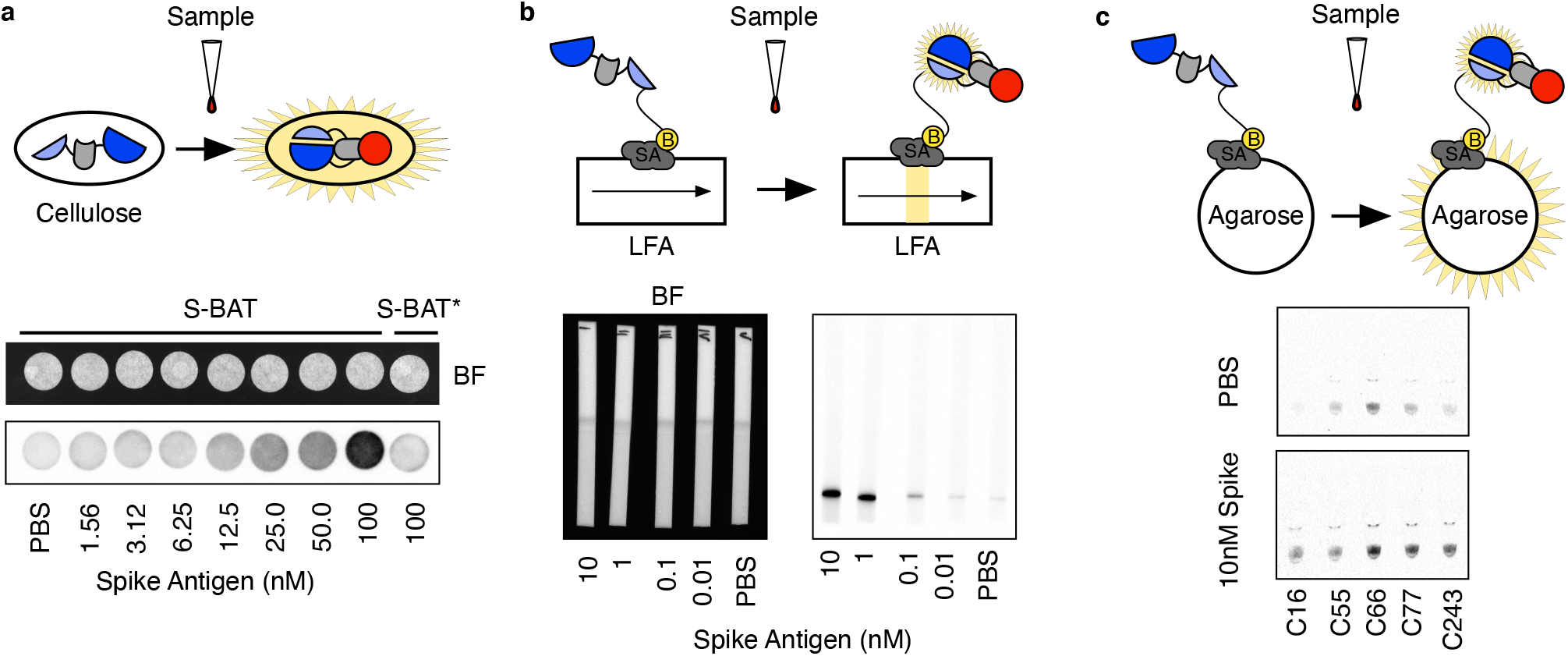
Application of S-BAT to immobilized assay formats. a) S-BAT or S-BAT* was adsorbed onto cellulose paper and dried in a vacuum desiccator. Spotting of antigen and substrate on the paper yielded detectable signal upon imaging with a CCD camera. The bright field (BF, top) and chemiluminescent (bottom) images for a serial dilution of recombinant Spike are shown. b) S-BAT (Cys16-PEG_11_-Biotin) was mixed with a serial dilution of recombinant Spike and substrate, and then resolved on a lateral flow assay (LFA) strip functionalized with streptavidin. The bright field (BF, left) and chemiluminescent (right) images demonstrate robust detection at 1 nM recombinant Spike concentration. c) Streptavidin functionalized agarose beads were functionalized with S-BAT (CysX-PEG_11_-Biotin) constructs and incubated with either buffer of 10 nM recombinant Spike. After addition of substrate, the beads were settled by centrifugation and imaged with a CCD camera. Biotinylation at Cys16 gave the best signal to noise ratio.

The lateral flow assay (LFA) is a workhorse platform for the rapid detection of biomolecules in clinical and laboratory settings.^20^ To demonstrate that BATs are amenable to use in the LFA format we screened our library of 10 conjugates (5 sites x 2 linker lengths) against 10 nM (1466.7 ng/mL) Spike. The sample (antigen or buffer), biotinylated sensor, and NanoGlo reagent were incubated in solution for 10 minutes. After incubation, a streptavidin functionalized LFA strip was immersed in the sample and allowed to incubate for an additional 10 minutes. The LFA strips were removed and imaged on a Bio-Rad ChemiDoc gel imaging system equipped with a CCD camera. The screen data demonstrated that the PEG_11_ constructs tended to yield higher signal in this format and that while conjugation at position 66 yielded the highest signal, immobilization via the N-terminus (position 16, P1’ of TEV cleavage site) gave the lowest background (**Figure S3a, b**). In this format S-BAT(Cys16-PEG_11_-Biotin) was able to detect 1 nM (146.7 ng/mL) recombinant Spike (**Figure 2b**).

The ELISA is a staple of clinical and laboratory testing. However, ELISAs require multiple functionalized antibody reagents and multiple rounds of washing and development steps. We envisioned using immobilized BATs to create an assay with the same form factor but that required no washing and fewer reagents. To this end, we immobilized S-BAT (Cys16-PEG_11_-Biotin) on a white 96-well plate pre-coated with streptavidin. Addition of sample and substrate to the well was sufficient to enable detection of low nanomolar concentrations of antigen by imaging without any washing steps in 10 minutes (**Figure S3c**). The LFA and plate assays are versatile assay formats; however, both are limited in the volume of sample that they can accommodate. To circumvent this limitation, we developed a bead-based reagent that could be incubated with a larger sample volume, isolated by centrifugation or filtration, and would concentrate signal from the sample at the time of measurement. We screened our library of biotinylated S-BAT constructs immobilized on streptavidin agarose beads against 10 nM (1466.7 ng/mL) recombinant Spike in reactions where the bead volume was 5% of the total volume. The beads were settled by centrifugation and analyzed by imaging the tube with a CCD camera. These results were consistent with our previous screen in the LFA format in that PEG_11_ linkers yielded the highest signal and conjugation at Cys16 yielded the highest signal to noise ratio (**Figure 2c**). BATs appear to be a flexible platform for protein analyte detection in multiple assay formats.

### BATs accommodate purification-free detection of antigen

While the production of protein reagents such as S-BAT in bacteria is simple and low-cost relative to the production of antibody reagents, it still requires significant protein purification infrastructure to achieve. We wanted to develop a protocol for the production and use of S-BAT that did not require purification. In a first iteration we simply expressed S-BAT and S-BAT* in the cytoplasm of BL21(DE3) *E. coli* with no affinity purification tag. A chemical lysis step followed by centrifugation and filtration yielded the clarified lysate as the working reagent. Dilution of the lysate and addition of antigen demonstrated that the crude lysate could detect recombinant Spike with sensitivity and specificity on par with the purified protein reagent (**Figure 3a**). Despite the low input necessary to culture *E. coli* at scale, certain aspects of their cultivation make them a suboptimal expression host screening S-BAT constructs. Moreover, while S-BAT was able to function in the context of crude cell lysates, a production system that did not require the chemical lysis was desirable. To this end we produced strains of *S. cerevisiae* that secreted S-BAT and S-BAT* into their growth media. We found that filtered culture media from these yeast were able to yield sufficient signal to detect 1 nM (146.7 ng/mL) recombinant Spike after 24 hours of induction of S-BAT expression (**Figure 3b**). These experiments demonstrate that S-BAT is robust in that it can function as a component of complex mixtures. In instances where the production of purified antigen is a challenge, such as with transmembrane protein targets, detection via BATs in intact mammalian cells streamlines the identification of active sensors. To show that BATs could function in this capacity we expressed full-length SARS-CoV-2 Spike, which contains a single-pass transmembrane anchor at its C-terminus, in HEK-293T cells. After 24 hours of expression we incubated cells with either 10 µM S-BAT or S-BAT* for 1 hour in complete cell culture media. The cells were washed and incubated with live cell Nano-Glo reagent in complete culture media. The cells were imaged with a CCD camera. These experiments demonstrated that signal was dependent on the expression of Spike and the integrity of LCB1 binding (**Figure 3c**). In total these experiments that BATs can facilitate the detection of antigen in crude formats.

**Figure 3.**
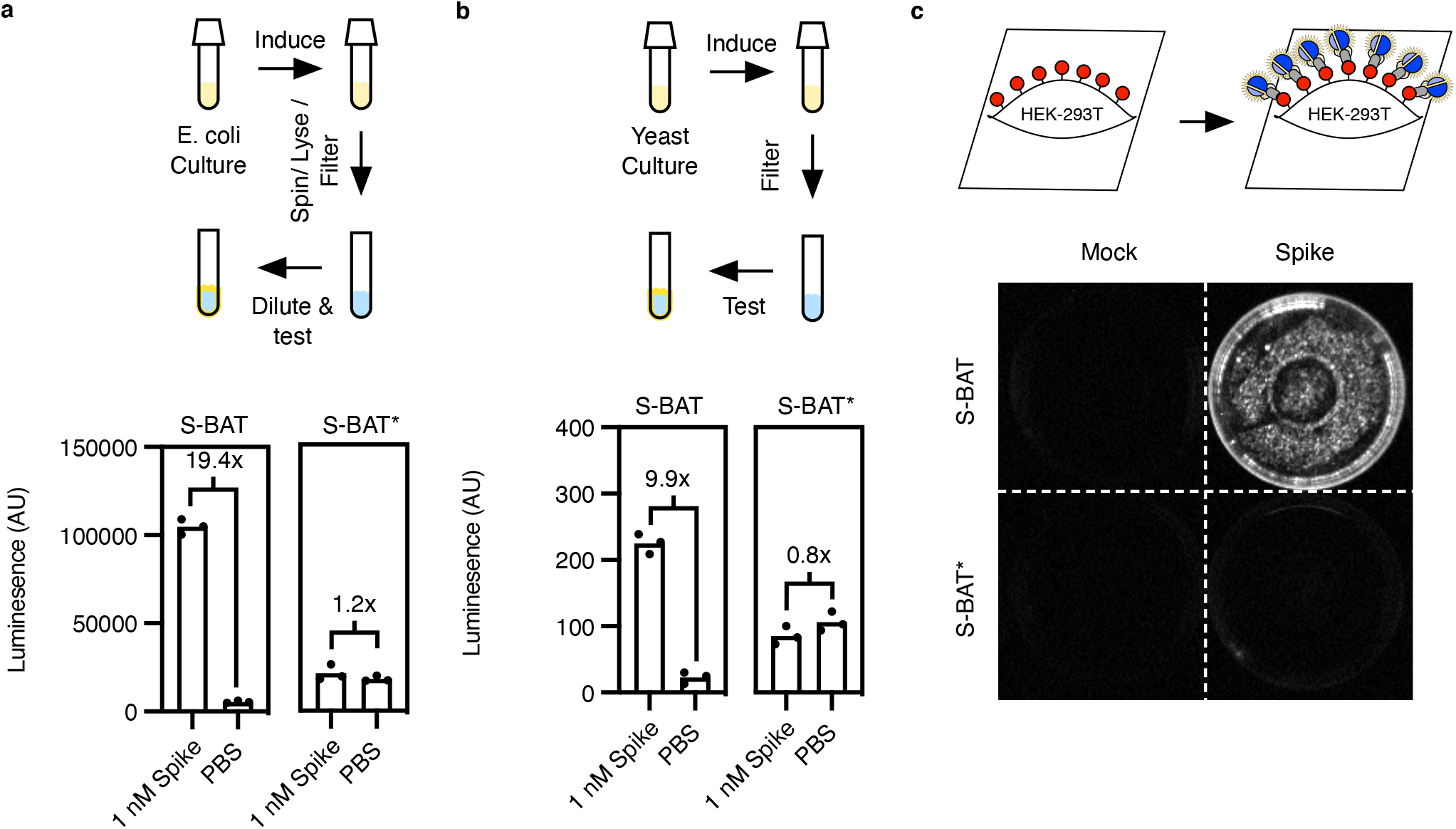
Purification-free production of S-BAT in bacteria and yeast. a) In a purification-free protocol for generation of S-BAT and S-BAT* in *E. coli*, the culture is induced with IPTG at room temperature, centrifuged, chemically lysed, and filtered to yield a crude extract. This extract is diluted and incubated with antigen prior to the addition of substrate. The diluted extract of S-BAT yields significant signal increase (19.4-fold increase in mean signal intensity) upon incubation with 1 nM recombinant Spike. In contrast, S-BAT* signal is not changed by addition of recombinant Spike. b) Yeast strains engineered to secrete S-BAT and S-BAT* are induced at 30 ºC for 24 hours, filtered, and the filtered media is used as a working reagent for detection without further processing. S-BAT produced in this manner yielded a 9.9-fold change in mean signal intensity upon incubation with 1 nM recombinant Spike, while S-BAT* showed no change in signal. Luminescence is plotted in arbitrary units (AU) for individual replicate samples (n=3) in a representative experiment. c) S-BAT, but not S-BAT* can specifically detect Spike expressed on the surface of HEK293T cells.

### Detection of cultured SARS-CoV-2

As the SARS-CoV-2 pandemic has continued, the emergence and characterization of dominant variants has been a significant focus of research efforts. With time and transmission, the sequence composition of the spike protein, and in particular the RBD have changed.^21^ LCB1 was originally designed to competitively antagonize the interaction between ACE2 and the RBD domain of the spike protein, the critical protein-protein interaction required for cell entry. Since this interface is critical to viral entry and propagation it might be expected that mutation to the interface would in general be deleterious to the virus.^22^ However, as the determinants of binding for ACE2 and LCB1 to RBD are overlapping but not identical, the probability that mutations could arise that differentially affect native receptor versus LCB1 binding is significant. We tested a panel of purified RBDs derived from sequences of prominent variants that have arisen throughout the pandemic (WA1, B1.1.7, B1.351, P1, B1.617.1 and B1.617.2) Given that B1.351 and P1 variants have a mutation at K417, which forms the key salt bridge with D30 in LCB1, we anticipated that these variants would completely evade detection by S-BAT. B1.1.7 contains a N501Y mutation in the RBD, which also has a high probability of disrupting S-BAT recognition (**Figure 4a**). Indeed, when screened against 100, 10, and 1 nM of each recombinant RBD, we were unable to detect B1.351 and P1 variants, while B1.1.7 showed an attenuated signal. Both B1.617.1 and B1.617.2 were readily detected at this concentration, consistent with the fact that they each carry RBD mutations distal to the ACE2/LCB1 binding site (**Figure 4b, Figure S4a**).

**Figure 4.**
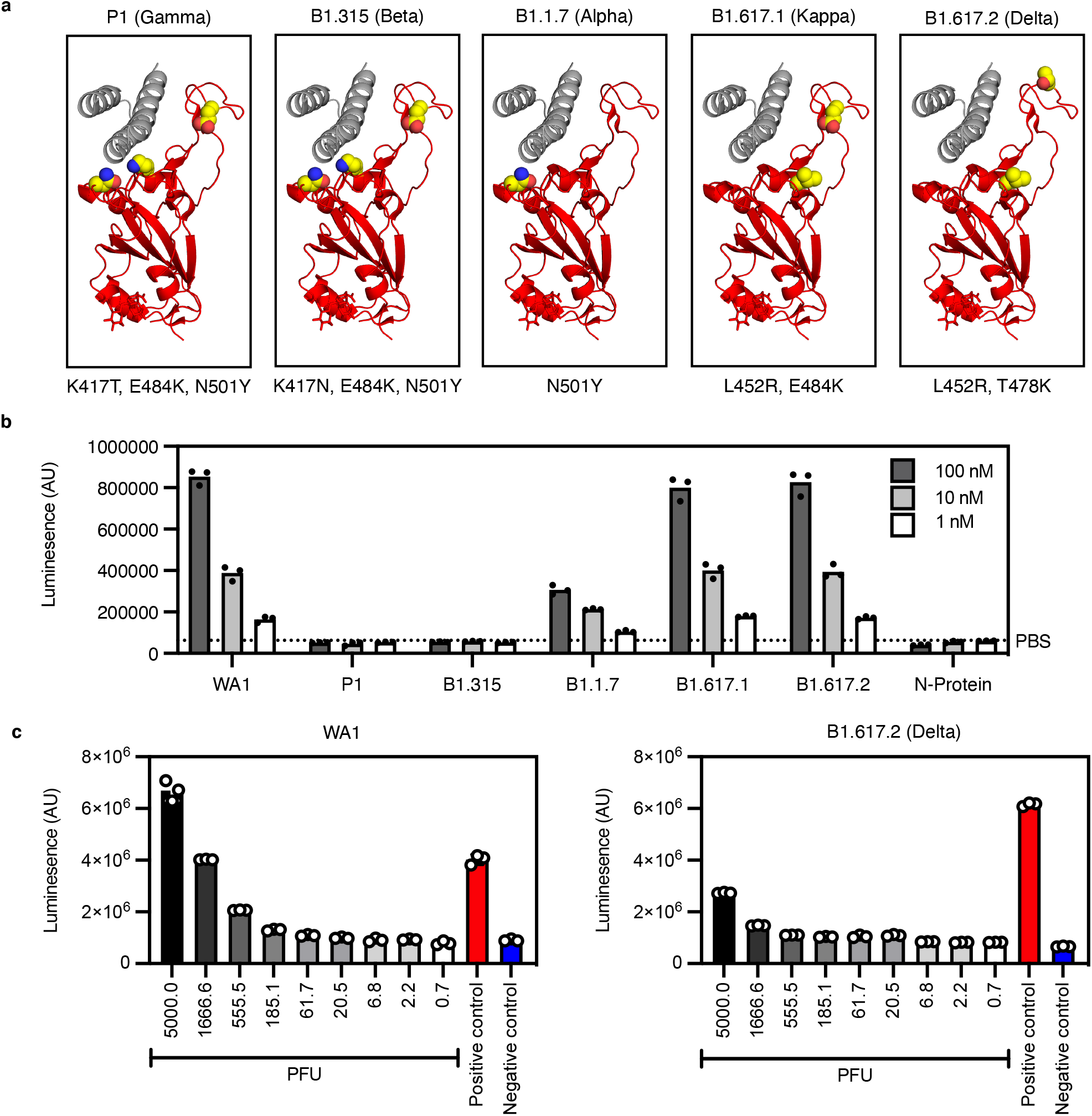
Detection of cultured SARS-CoV-2 variants with S-BAT. a) The positions of receptor binding domain (RBD, red, cartoon) mutations (yellow, sphere) in variants of concern relative to the LCB1 (grey, cartoon) binding site. b) Screening of recombinant RBDs at 100, 10, and 1 nM from variants of concern WA1, P1 (Gamma), B1.1.7 (Alpha), B1.315 (Beta), P1.617.1 (Kappa), and P1.617.2 (Delta) predicted that WA1, Kappa, and Delta would be robustly detected by S-BAT. N-Protein yielded no signal in the assay. c) S-BAT detection of live cultured virus demonstrates the predictive power of the recombinant antigen screen. S-BAT successfully detected both WA1 and Delta variants of SARS-CoV-2. Mature virus particles diluted serially in media (greyscale bars) were compared to 1 nM recombinant Spike (red, bar) and media (blue, bar). Virus was tittered and standardized by plaque assay and diluted based of plaque forming units (PFU). Luminescence is plotted in arbitrary units (AU) for individual replicate samples (n=3) in a representative experiment.

To understand the correlation between our in vitro RBD assay and our ability to detect cultured, live SARS-CoV-2 virus, we selected WA1 and B1.617.2 (Delta) as probable positive variants and B1.351(Beta) as a probable negative variant. Virus was screened at concentrations consistent with clinical isolates from nasopharyngeal swabs.^23^ As predicted by the results with recombinant protein, we were able to detect cultured virus from WA1 (the first strain detected in the United States) and B1.617.2 (Delta variant) and unable to detect cultured virus from B1.351(Beta variant, first detected in South Africa) (**Figure 4c, Figure S4b**). S-BAT* signal did not change when incubated with any cultured virus variant (**Figure S4b, c, d)**. The predictive value of the recombinant antigen assay for cultured virus detection lowers the activation barrier for testing and adapting this and any future generations of S-BAT against variants of concern as they emerge.

### Cell phone detection of antigen

Beyond making a reagent that is easy to produce, we wanted to develop a detection platform that takes advantage of the ability of S-BAT to generate visible light, and the fact that most people walk around with a moderately sensitive photodetector in their pocket in the form of a cell phone camera. The key technical hurdles in detecting S-BAT activation with a cell phone camera are sufficiently eliminating ambient light and maximizing photon collection from the sample. To achieve this, we designed and fabricated an adaptor to an iPhone SE (2020) with the form factor of a cell phone case (**Figure 5a**). The case was designed to accommodate a homogenous sample placed in a disposable microcuvette (**Figure 5b**). The case itself was fabricated on a 3D printer from black ABS thermoplastic based on its high opacity. The inside of the reaction chamber was covered with a diffuse reflective coating to minimize the loss of emitted photons.^24^ The cuvette was positioned as close as possible to the camera aperture. RAW images were acquired, and a region of interest (ROI) was selected based on where in the image the blue light intensity was highest. The use of an optimized ROI rather than the entire image enhanced the dynamic range of the measurement (**Figure 5c, Figure S5**). To define a limit of detection (LOD) for the cell phone acquisition we incubated decreasing amounts of WA1 recombinant Spike with 10 nM S-BAT and compared the mean intensity to vehicle (PBS buffer). S-BAT* incubated with recombinant Spike and S-BAT incubated with BSA yielded no detectable increases in signal (**Figure 5d, e**). Using a cutoff of 3***σ*** we were able to detect a lower limit of 300 pM (44.0 ng/mL) recombinant Spike with this detection method. This sensitivity is on par with the reported molar LOD for commercial antigen test sensitivities for recombinant antigen.^25^ The pairing of S-BAT and cell phone detection of luminescence are a promising platform for in-field analyte detection.

**Figure 5.**
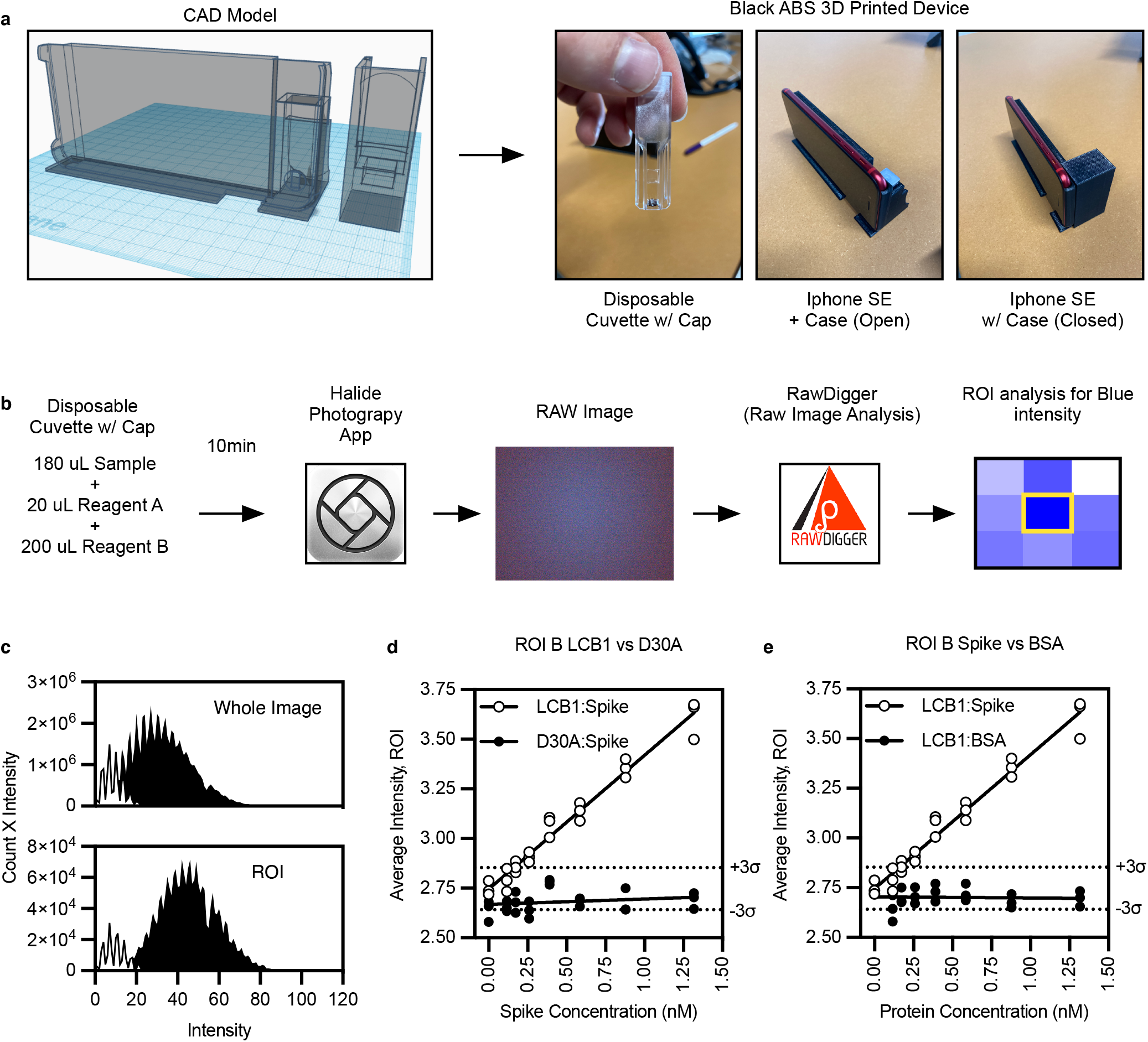
Detection of SARS-CoV-2 S-protein using S-BAT and a cell phone. a) Computer-aided design (CAD) model of cell phone case (left) and cover (right) for low-light imaging of chemiluminescence. b) Reactions were set up in a disposable microcuvette with a cap (left panel), placed into the iPhone SE cell phone case 3D printed in ABS thermoplastic (center panel), and capped to exclude ambient light (right panel). c) Reactions were set up in 400 µL with 180 µL sample, 20 µL of 200 nM S-BAT or S-BAT*, and 200 µL 2X substrate, incubated for 10 minutes, and a RAW image was taken using the HALIDE photography application. Since the signal in the image was not homogenous, a grid search of a prototype image was used to identify a region of interest (ROI) with the highest signal intensity using the RAW image analysis software RAWDigger. d) Use of the optimized ROI (bottom histogram) rather than the entire image (top histogram) yielded better separation between a high signal sample (100 nM recombinant Spike, black) and negative control (PBS, white) Data are plotted as pixel intensity x pixel count such that the area under the histogram represents the total blue intensity for the analyzed region. e) Plots of average blue intensity for ROI of S-BAT incubated with recombinant Spike (open circle, left panel) or S-BAT* incubated with recombinant Spike (filled circle, left panel). The same plot for S-BAT incubated with recombinant Spike (open circle, right panel) or S-BAT incubated with bovine serum albumin (BSA) (filled circle, right panel). Replicates are three images taken of the same sample in sequence. Solid lines are a linear regression of each data set. Dashed lines are ±3**σ**of the negative control (PBS). Average Intensity is plotted in arbitrary units (AU) versus antigen concentration (nM) for technical replicates (n=3) in a representative experiment.

## Discussion

In the specific context of the SARS-CoV-2 pandemic, S-BAT is a potentially useful reagent for assaying cultured virus, pseudo-virus, and components of subunit vaccine candidates. The limits of detection we reported here may be sufficient to detect Spike antigen at concentrations reported in urine and plasma from patient samples by sandwich ELISA.^26,27^ While the BAT platform itself is promising for diagnostic development, the S-protein may not be as well-suited for SARS-CoV-2 detection as the N-protein, which is much more abundant and accessible in patient samples despite its putative localization in the intact virion. Moreover, S-protein accumulates more mutations than N-protein in emerging variants. This liability is exemplified by the emergence of the omicron variant which is mutated relative to the WA1 ancestral strain at K417 and N501. Based on the ablation of activity by K417 mutation in the beta and gamma variants, and the attenuation of activity by the N501 mutation in the alpha variant, we would anticipate that S-BAT would not detect the omicron variant.

We have demonstrated the versatility and sensitivity of the BAT platform in multiple assay formats. This use case, the detection of SARS-CoV-2 protein antigen, highlights the importance of attributes beyond specificity and sensitivity that should be considered when developing new technologies. BATs are a first step towards a reagent class that is amenable to being produced and used with relatively low-cost infrastructure. Despite the meaningful advantages that BATs possess relative to antibody-based detection methods in this sense, significant improvements can be made to the platform to enhance its utility. While the NanoBiT system is convenient and well-studied for the purposes of developing biosensors, the small molecule luciferin substrate it uses, Furimazine, is expensive, unstable, and proprietary. Adapting BATs to incorporate a different reporter enzyme system such as peroxidases, alkaline phosphatase, or other non-proprietary luciferase enzymes may yield systems that can incorporate more robust and accessible substrate classes with the capacity to yield both luminescent and colorimetric readouts. Future work will focus on adapting BATs to other antigens and further exploring the modularity of this platform.

## Materials and Methods

### Cloning

All BAT constructs were ordered from Twist Biosciences (See DNA and Protein sequences in Supplemental Information) and were subcloned into a pET backbone (https://www.addgene.org/29659/) using NcoI and BamHI restriction sites via Gibson assembly. For yeast secretion constructs, sequences for S-BAT and S-BAT* were subcloned into the pCTCON2 backbone (https://www.addgene.org/73152/) using the EcoRI and XhoI restriction sites via Gibson assembly.

### Recombinant Protein Production

Plasmids encoding BAT sensors were used to transform E. coli BL21 (DE3) competent cells (Agilent). Transformants from a single colony were used to inoculate 5 mL of LB supplemented with 100 mg/L kanamycin (LB-Kan) and grown overnight at 37 ºC, 220 rpm. The overnight seed culture was diluted 1:100 LB-Kan medium and was incubated at 37 ºC, 220 rpm until the OD600 was between 0.4 and 0.6. The cells were then induced with by adding IPTG to a final concentration of 1 mM and grown for another 20 hours at room temperature. After induction, cells were harvested by centrifugation at 10,000 g for 20 minutes. The cell pellet was resuspended in 1:4 w/v (g/mL) B-PER Reagent (Thermo Fischer) supplemented with PMSF and Protease Inhibitor Cocktail and incubated for 15 minutes at room temperature. The cellular debris was removed with centrifugation (21,300 g, 30 minutes, 4 ºC), and the supernatant was cleared using a syringe filter (0.45 µm, PVDF, Millipore). The filtered lysate was batch loaded to 0.5 mL of packed Ni-NTA resin. The column was washed 2 times with 5 mL wash buffer (50 mM Tris-HCl pH 8.0, 300 mM NaCl, 30 mM Imidazole). The protein was then eluted using for a total of 4 mL elution buffer (50 mM Tris-HCl pH 8.0, 300 mM NaCl, 200 mM Imidazole). The fractions were analyzed using SDS-PAGE, and the fractions containing the un-cleaved BAT sensor were collected. The un-cleaved BAT sensors were concentrated to ∼1.0 mL, and the concentration was calculated from the A280 value. The His_6_ tag at the N-terminus was cleaved by adding the His-tagged TEV protease at 1:50 molar ratio to the protein. The mixture was dialyzed into TEV reaction buffer (20 mM Tris-HCl pH 8.0, 150 mM NaCl, 0.5 mM TCEP-HCl) overnight at 4 ºC. The mixture was loaded to Ni-NTA resin, and the cleaved BAT sensor was collected by keeping the flow-through. The cleaved BAT sensor was dialyzed into PBS pH 7.4 overnight at 4 ºC. The purity of the cleaved sensor was analyzed using SDS-PAGE, and the final product was stored 4 ºC for further use. S-BAT and S-BAT* were further purified by size exclusion chromatography on a HiLoad 16/600 Superdex S75 column run on an Akta go protein purification system (Cytiva) in PBS. S-BAT variants containing single cysteine residues were purified by the same protocol but dialyzed into 20 mM HEPES pH 7.4, 150 mM NaCl, 1 mM TCEP. Biotin conjugates were prepared by the addition of 2 equivalents of Maleimide-PEG_2_-Biotin or Maleimide-PEG_2_-Biotin overnight at 4 ºC. Excess biotinylation reagent was removed by iterative dilution and concentration in PBS pH 7.4. Tag-free S-BAT produced in E coli were expressed and extracted as above on a 5 mL scale. After chemical lysis, the crude mixture was filtered through a 0.45 µm PVDF syringe filter. The filtrate was diluted 1:100 in PBS pH 7.4 prior to use.

The pCTCON2 plasmids encoding the secreted BAT sensors were transformed into S. cerevisiae strain EBY100 using Frozen-EZ yeast transformation Kit II (Zymo Research). Transformants from a single colony on Trp-deficient SDCAA plates were grown and used for subsequent yeast experiments. Trp-deficient SDCAA and SGCAA media were used, for propagating and inducing yeast cells harboring the S-BAT sensor secretion plasmid, respectively. Yeast cells were induced by inoculating 1.8 mL of SGCAA medium with 0.2 mL of saturated yeast culture (OD600 = 15) maintained in SDCAA medium. After inducing for 24 hours at 30 ºC, 250 rpm, the cells were pelleted by centrifugation at 3,000 g for 2 minutes. The supernatant was cleared using a syringe filter and used for subsequent detection assay.

Plasmids encoding SARS-CoV-2 Spike (S) ectodomain or Receptor Binding Domain (RBD) were transiently transfected into suspension Expi293 cells at 1-1.5 L scale in 2.8 L Optimum Growth Flasks (Thomson Scientific), following the manufacturer’s guidelines. Three days after transfection, cell cultures at >75% viability were centrifuged at 500 g for 30 min, followed by filtration of the supernatant using a 0.45 µm Nalgene Rapid Flow filter unit. The supernatant was adjusted to pH 7.4, and directly loaded onto a 5 mL HisTrap Excel column pre-equilibrated with 20 mM sodium phosphate, 500 mM NaCl, pH 7.4 using an AKTA Pure. Captured proteins were washed with 60 column volume (CV) of (20 mM sodium phosphate, 500 mM NaCl, 20 mM imidazole, pH 7.4) and eluted with 10 CV of (20 mM sodium phosphate, 500 mM NaCl, 500 mM imidazole, pH 7.4). Eluted proteins were buffer exchanged into PBS using either 3kDa MWCO (for RBD) or 100 kDa MWCO (for Spike) Amicon concentrators, supplemented with 10% glycerol, and filtered prior to storage at -80 °C.

RBD’s from SARS-CoV2 variants of concern and N-Protein were purchased from Sino Biological, and reconstituted from lyophilized powder as per the manufacturer’s instructions.

### Homogenous detection of recombinant Spike or RBD

All BAT sensors were diluted into PBS pH 7.4 to the final concentration of 4 nM prior to the Spike detection assay. The final concentration of sensor in these assays was 1 nM. The stock of recombinant Spike was diluted into PBS pH 7.4. 20 μL of diluted sensor was mixed with 20 μL of diluted recombinant Spike in a PCR tube, mixed well by pipetting, and incubated for 1 hour at room temperature. 25 μL of this mixture was mixed with 25 μL of Nano-Glo® Luciferase Assay Reagent (Promega), following manufacturer’s protocols, in a white, opaque 384-well plate. The solution was further mixed by shaking, incubated for 10 minutes at room temperature, and the luminescence was measured using a plate reader (M1000, TECAN). For purification free generation of S-BAT, *E. coli* lysate was diluted 1:100 in PBS pH 7.4 prior to use in this assay. For S-BAT secreted from *S. cerevisiae*, and filtered culture media was used as is in place of the purified S-BAT.

### Cellulose Paper Assay

To adsorb S-BAT onto cellulose paper one 10 mm diameter round cellulose filter paper (Whatman Grade 1) was placed in a 48 well polystyrene tissue culture plate to ease handling and batch production. The paper was submerged in a solution of 10 mg/mL bovine serum albumin (BSA) and 0.2% Tween-20 for 1 hour at room temperature. The blocking solution was removed and 100 µL of 100 nM S-BAT in PBS pH 7.4 was added to the well for 1 hour. The S-BAT solution was removed, and the entire plate was placed in a vacuum desiccator for 48 hours. Papers were removed and stored in a conical tube at room temperature until use. The detection assay was conducted by mixing sample and 2X (1:50 dilution) NanoGlo reagent in NanoGlo buffer and spotting 25 µL on the paper. After 10 minutes the paper was imaged using a ChemiDoc XRS (Bio-Rad) imaging system with a 1 s exposure time.

### Lateral Flow Assay

Lateral flow assays (LFA) were conducted using off-the-shelf streptavidin functionalized test strips (GenLine, Milenia Biotec) which were modified by removing the sample pad. A 40 µL reaction was set up using 10 µL of 40 nM biotinylated S-BAT, 10 µL of sample, and 20 µL of 2X NanoGlo reagent. The reaction was incubated for 10 minutes and the LFA strip was placed in the reaction and allowed to develop for 10 minutes Test strips were removed from the reactions and imaged using the same protocol as the Cellulose paper assay.

### Bead Assay

Neutravidin functionalized agarose beads were washed in PBS pH 7.4 loaded with 100 nM biotinylated S-BAT for 1 hour at 4 ºC in a 50% slurry. After loading beads were washed three times with 40 bead volumes of PBS pH 7.4 and stored as a 2.5% slurry. For detection assay 50 µL of 2.5% slurry was mixed with 50 µL of sample for 10 minutes in a thin-walled PCR tube. 100 µL of 2X NanoGlo reagent was added, and the beads were pelleted by centrifugation. Pelleted beads were imaged in the tube using the same protocol as the cellulose paper and LFA assays.

### Live-cell detection of SARS-CoV-2 Spike protein

HEK293T cells were seeded in a 35 mm tissue culture dish coated with 1 mg/mL human fibronectin. 7.5 × 10^5^ cells were plated and grown overnight in complete DMEM (10% fetal bovine serum (FBS), 1% penicillin/streptomycin). The following day the cells were transfected with 500 ng of plasmid DNA encoding the full length WA1 SARS-CoV-2 spike protein behind a CMV promoter using PEI. After 5 hours the media was changed, and cells grown for an additional 20 hours. S-BAT and S-BAT* were diluted to 10 nM in complete media and incubated with cells for 1 hour. Media was removed, and cells were washed 3X with complete media. Live cell NanoGlo (Promega) was diluted in complete media and 1 mL was added to the cells. Cells were imaged on a ChemiDoc XRS (Bio-Rad) imaging system with a 10 s exposure time.

### SARS-CoV-2 Virus culture and detection

All SARS-CoV-2 live virus experiments were performed in a biosafety level 3 facility with approved procedures at the Gladstone Institutes. SARS-CoV-2 isolates and variants USA-WA1/2020, B.1.351 (beta), and B.1.617.2 (delta) were procured from BEI resources. The virus was cultured in Vero-E6 cells (obtained from ATCC and grown in DMEM (Corning) supplemented with 10% fetal bovine serum (FBS) (GeminiBio), 1% glutamine (Corning), and 1% penicillin-streptomycin (Corning) at 37 °C, 5% CO2) and stocks were stored at -80°C. Virus titer was determined via plaque assays on Vero-E6 cells. Briefly, 0.2 × 10^6^ cells per well were seeded in 12 well plates, incubated overnight, and infected with a range of virus dilutions (10^1^-10^6^) for 1 hour prior to overlay by 2.5% Avicel (Dupont, RC-591. After 72 hours, the Avicel was removed, cells were fixed in 10% formalin for 1 hour and stained with crystal violet for 10 minutes to visualize plaque forming units.

For S-BAT detection experiments, 25 µl of virus dilutions were combined with 25 µl of Spike sensor (diluted in PBS from 4X stock) in a 96-well white polystyrene microplates (Corning) and incubated for 1 hour at room temperature. Next, 50 μL of 2x Nano-Glo buffer was added and mixed by gentle pipetting, the plate was sealed and incubated at room temperature for 10 minutes, and luminescence was measured (Promega Glowmax Luminometer). Recombinant Spike and media served as positive and negative controls, respectively.

### Cell Phone Detection

The cell phone adaptor was designed to fit an iPhone SE (2020) and accommodate a disposable semi-microcuvette (FisherScientific Cat No. 14955127) using the Tinkercad design platform (tinkercad.com). The adaptor was printed as two parts on an FlashForge Creator Pro 2 3D printer using black ABS filament. The interior of the reaction chamber was coated with an adhesive diffuse reflective coating (BrightWhite 98, Thin Film with Adhesive and Liner, R-MG98-xx06-AD01) to enhance signal retention. Reactions were set up by combining 180 µL of sample with 20 µL of 200 nM S-BAT or S-BAT* and 200 µL of 2X NanoGlo reagent in the cuvette. The reactions were incubated for 10 minutes prior to image acquisition. RAW Images were acquired with the Halide photography application (Focus +1, ISO 2122, S1/3) and exported to RAWDigger for analysis. A region of interest (ROI) was defined to maximize signal in the blue channel using a template image. The ROI was chosen based on a grid search for peak intensity. Briefly the image was divided into 16 even regions and the average blue intensity for each was calculated. This process was repeated within the region of highest signal. The maximal signal intensity was detected in a 252×189 pixel region starting at (x, y) coordinates of (2016,1512). This region was used for the analysis of all samples.

## Author Contributions

M.R. and H.R. contributed equally. Project conception M.R. and A.Y.T. Protein production M.R. H.R and J.P. Assay development M.R. and H.R. Cell phone detection platform M.R. Live virus BSL3 studies R.S., G.R.K., and M.O. Writing M.R. H.R. and A.Y.T. Supervision A.Y.T. and M.O. Editing/Review all authors.

## Notes

The authors declare no competing financial interests

## Acknowledgments

This work was supported by funds from a Fast Grant (https://fastgrants.org/) and a generous donation from Jonathan Levett and the Marin Community foundation. M.O. received support from the Roddenberry Foundation and a gift from Pam and Ed Taft. M.R. was supported in part by a training grant from the National Human Genome Research Institute (NHGRI 5T32HG000044-24). The authors would like to thank Joseph L. Derisi (UCSF, CZ Biohub), James A. Wells (UCSF), Susanna K. Elledge (UCSF), Daniel A. Fletcher (UC Berkeley), Paul Lebel (CZ Biohub), and Rafael Gómez-Sjöberg (CZ Biohub) for input at various stages of this work. This work is dedicated to the memory of M.R.’s father-in-law, Michael B. Richardson, a committed science educator and a perpetually curious mind.

