## Supporting Info for "A single-component luminescent biosensor for the SARS-CoV-2 spike protein"

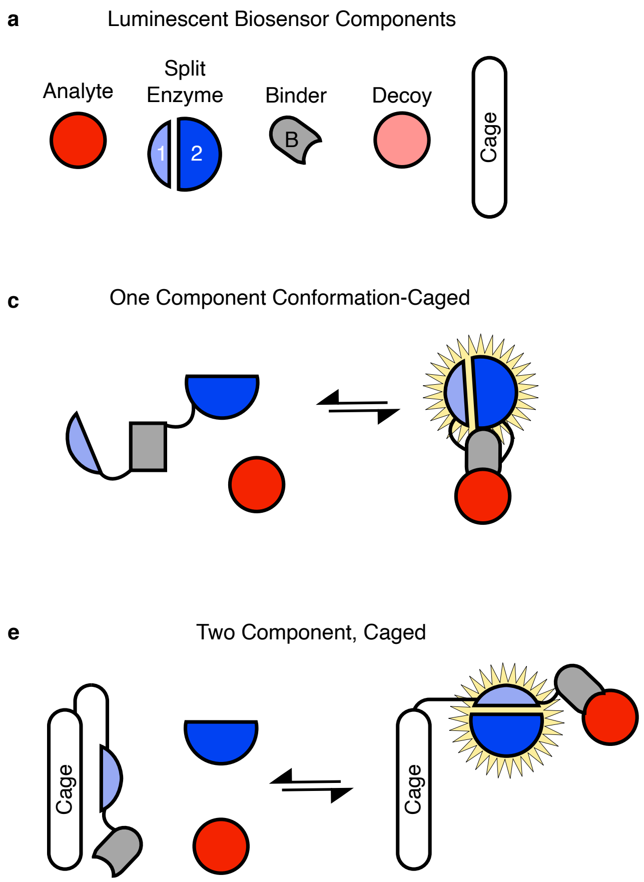


**Figure S1.** Chemiluminescent biosensor platforms. a) Existing platforms to detect an analyte of interest (red) generally contain of a split enzyme (1 & 2, Blue), a binding module (B, grey), and may contain a tethered decoy analyte (light red) or a caging structure (white). b) Two component systems rely on two binding modules to reconstitute the halves of the split enzyme. c) One component systems can produce specific signal in response to analyte by undergoing a conformational change in response to binding or d) by tethering a decoy analyte to the sensor that prevents activation until displaced by analyte. e) Caged two component systems rely on an engineered molecular switch that unmasks one half of the split enzyme upon binding of analyte to a single binding module, allowing it to interact with the complementary half of the split enzyme. f) Binding activated tandem split-enzymes (BAT) biosensors rely on the steric clashes between the bound analyte and the split enzyme components to drive sensor activation.

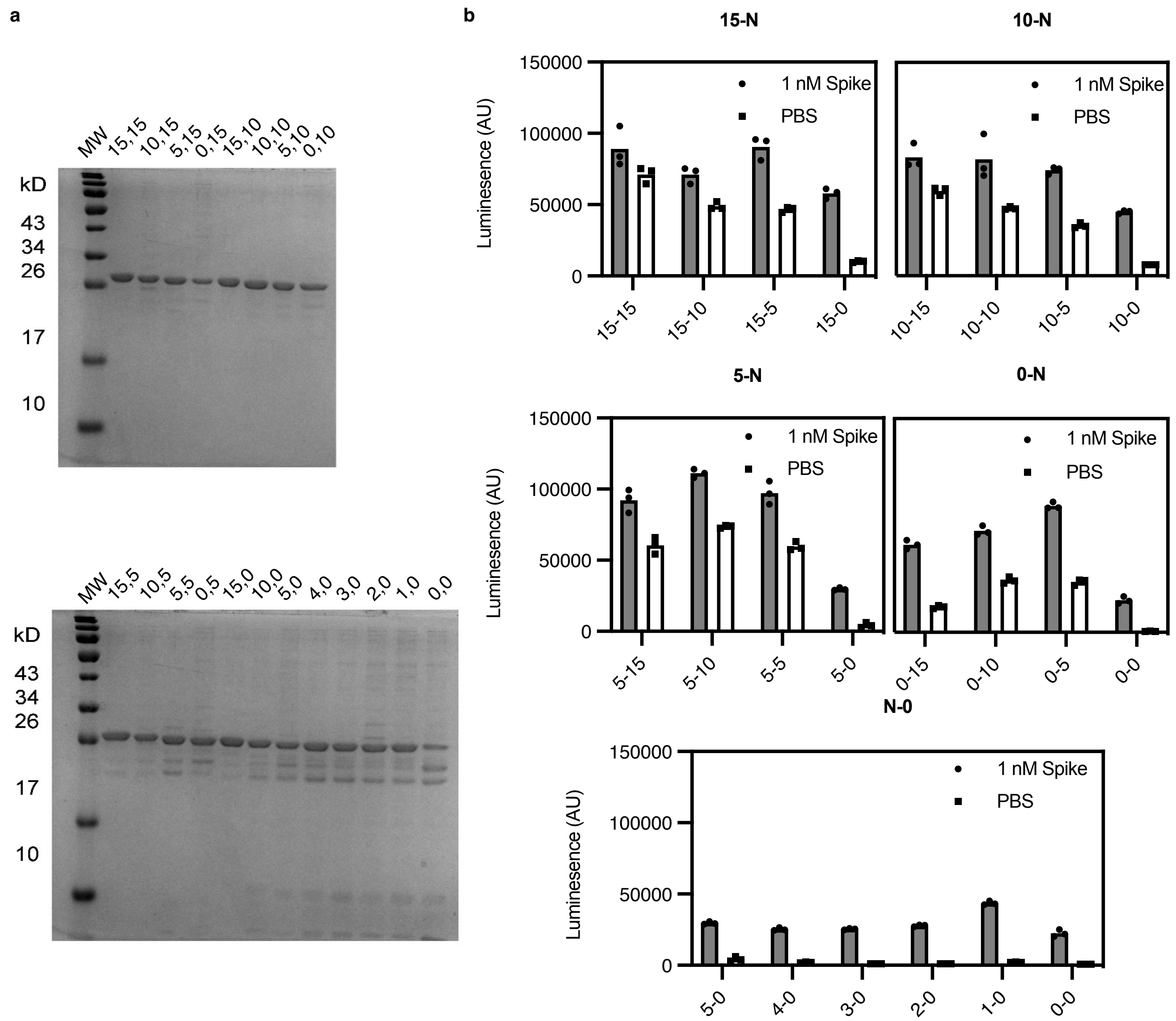


**Figure S2.** a) Analysis of linker library purity after histidine tag removal by TEV protease cleavage and subtractive Ni-NTA IMAC. 15% Tris-Glycine SDS-PAGE gel was resolved and visualized by Coomassie stain. A notable increase in degradation products was evident in constructs with shorter linkers. For 0,0 a significant portion of isolated protein was degraded. b) Raw data from the linker library screen used to generate the heat map in Figure 1c. Luminescence is plotted in arbitrary units (AU) for individual replicate samples (n=3) in a representative experiment.

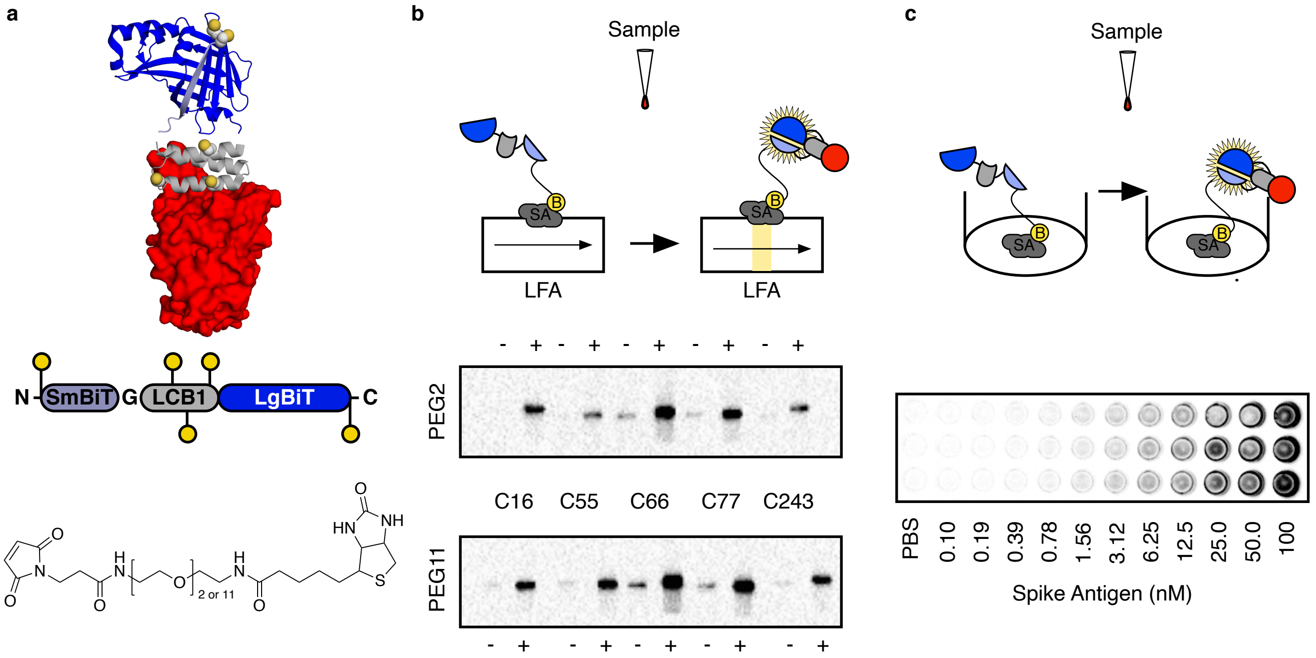


**Figure S3.** a) 5 different conjugation sites (C16, C55, C66, C77, and C243) and 2 PEG linker lengths (2,11) were selected to minimize perturbations to binding and activation of S-BATs. B) A screen of linker lengths and conjugation sites with PBS (-) and 10 nM recombinant Spike (+) led to the selection of Cys16-PEG$_{11}$-Biotin) construct for further experiments based on signal intensity and low background in a 1 s exposure on a ChemiDoc XRS imaging system. c) A streptavidin coated 96 well plate was coated with 100 nM S-BAT (Cys16-PEG$_{11}$-Biotin) in PBS. Antigen and NanoGlo reagent were added to the well in triplicate and incubated for 10 minutes. The plate was imaged on a ChemiDoc XRS imaging system (Biorad) using a 10 s exposure time.

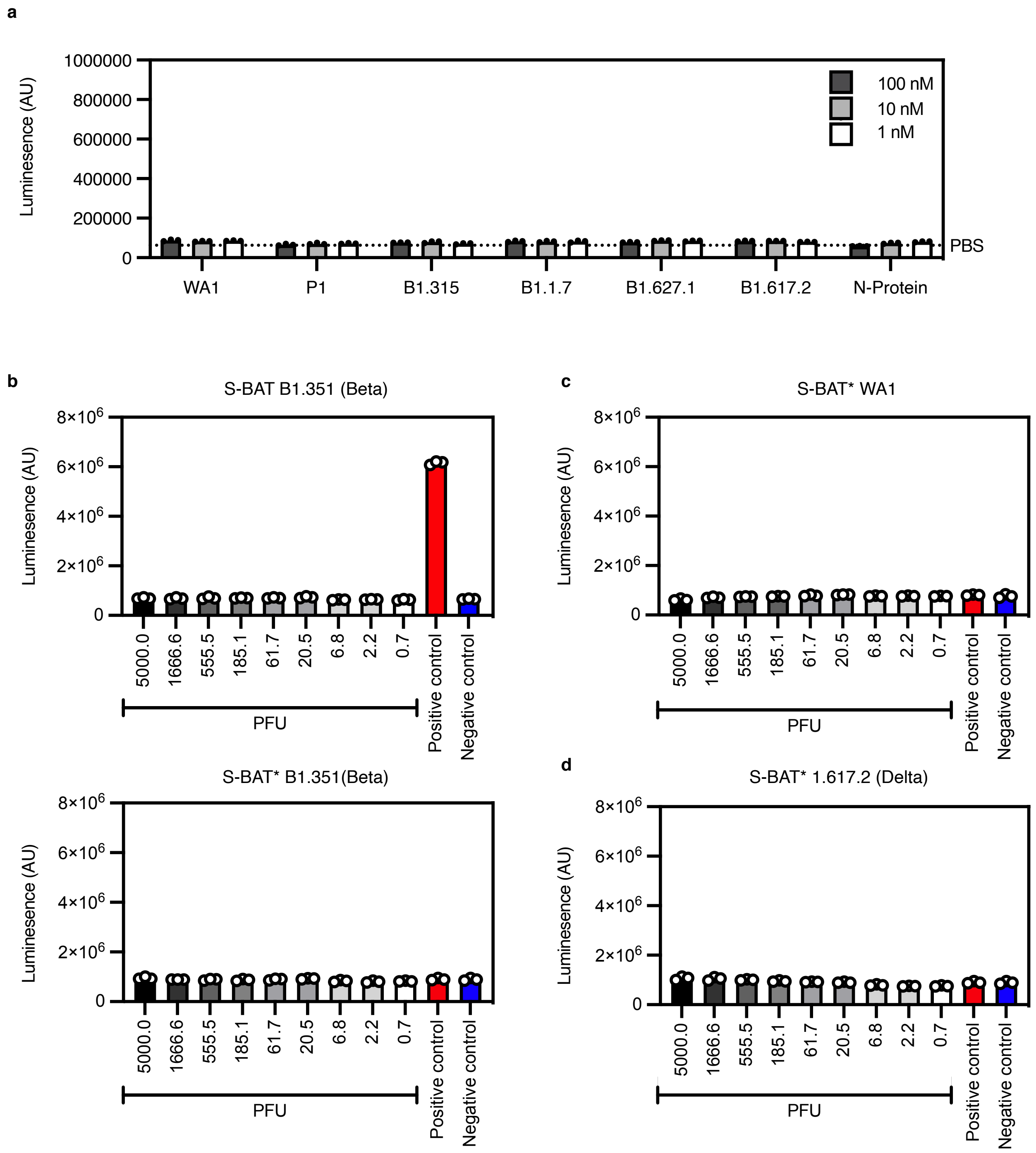


**Figure S4.** Negative controls for variant RBD and cultured virus detection. a) S-BAT* showed no change in signal relative to PBS for any variant RBDs or N-Protein at 100, 10, or 1 nM. b) Titration of cultured SARS-CoV-2 variant B1.351 (Beta) with the S-BAT sensor showed no activation above negative control (complete DMEM, blue) while positive control (1 nM recombinant WA1 Spike protein, red) activated the sensor. The same lack of activation was observed for S-BAT*. These data are in line with predictions from recombinant RBD experiments (Figure 4b). c,d) S-BAT* was not activated by WA1 or B1.617.2 (Delta) cultured virus supporting the specificity of the signal seen with S-BAT in Figure 4c. Luminescence is plotted in arbitrary units (AU) for individual replicate samples (n=3) in a representative experiment.

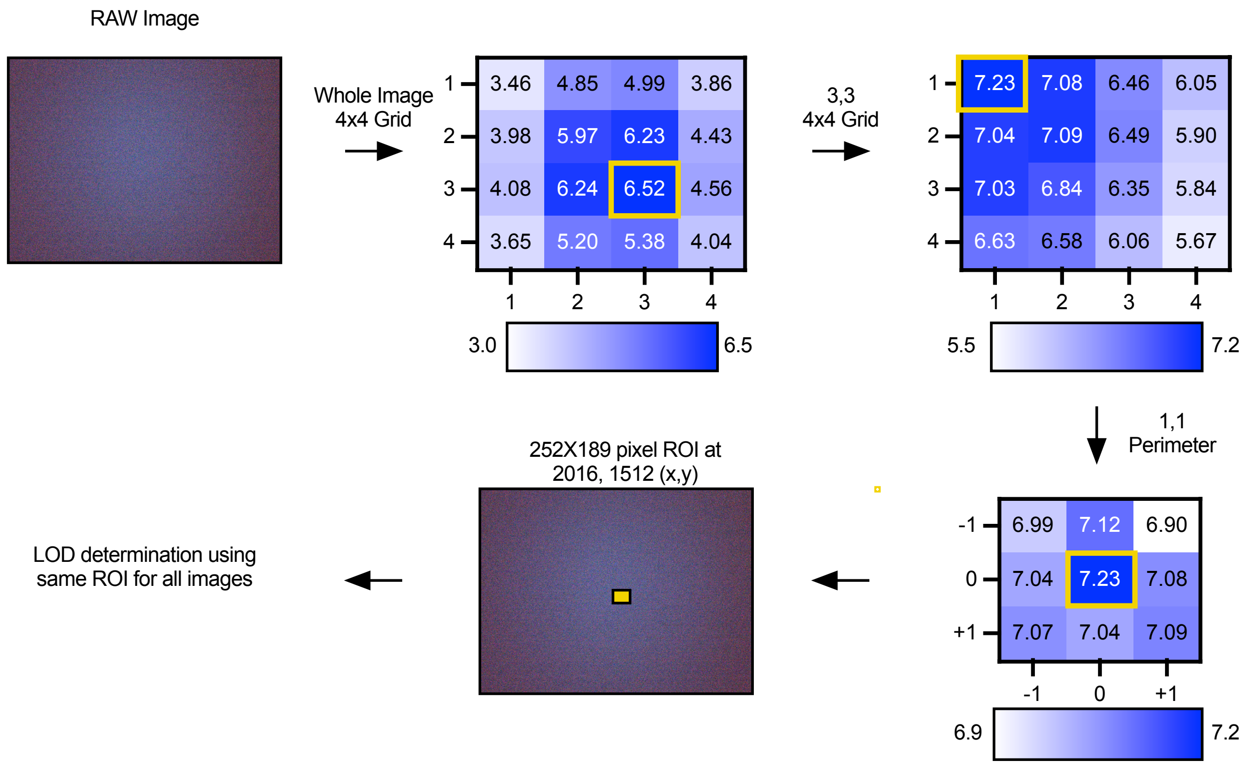


**Figure S5.** Grid search for ROI in RAW image from cell phone with maximum blue signal. Despite the samples being homogenous, images taken with the cell phone were visibly darker around the edges and brighter in the middle. To select an ROI with maximal signal in an unbiased fashion, a template image was broken into sixteen identically sized regions. The region with the highest average blue intensity (3,3) was broken into sixteen identically sized regions. The region surrounding with the highest average blue intensity (1,1) was then searched to ensure that a maximum had been identified. This ROI was used to analyze all samples for determining an LOD for Spike detection with S-BAT using the cell phone imaging protocol.

**DNA AND PROTEIN SEQUENCES**

**6xHis-TEVcs-SmBiT-15aa-LCB1-15aa-LgBiT**

ATGGCGCATCACCACCACCACCACGAGAATCTTTATTTCCAAGGTTCGGTGACCGGGTACCGTTTGTTCGAGGAGAT
TCTGGGTTCTGGATCAGGAGGGTCTGGAAGCGGCGGCTCAGGTTCCGGAGATAAGGAATGGATTTTGCAAAAGATC
TATGAGATTATGCGTTTACTTGACGAACTTGGACACGCTGAGGCGTCGATGCGTGTGTCGGACCTCATTTATGAGTTC
ATGAAGAAGGGCGACGAGCGCTTGTTGGAAGAGGCCGAGCGTCTCCTCGAAGAGGTCGAGCGCGGTTCTGGATCA
GGAGGTTCAGGAAGCGGCGGCAGCGGCTCAGGGATGGTATTTACGTTGGAAGATTTCGTGGGAGACTGGGAGCAA
ACCGCGGCCTATAACTTAGATCAGGTGTTGGAGCAAGGTGGTGTTTCGAGTTTACTGCAGAACCTTGCCGTGTCCGT
CACACCTATTCAGCGTATCGTCCGTTCAGGTGAGAATGCACTCAAGATTGACATTCACGTCATCATCCCGTACGAGG
GGTTAAGCGCAGACCAAATGGCTCAAATCGAAGAGGTATTTAAGGTCGTGTATCCAGTCGATGACCACCATTTTAAAG
TCATCCTGCCTTATGGCACATTGGTGATTGACGGGGTTACACCAAATATGTTGAACTATTTTGGACGTCCTTACGAGG
GGATCGCCGTGTTCGATGGAAAGAAGATCACGGTCACGGGGACGTTATGGAATGGTAACAAAATCATCGATGAGCGT
TTAATTACGCCGGACGGCTCGATGTTATTCCGTGTTACAATCAATAGCTGA
Corresponding to:
MAHHHHHHENLYFQ↓GSVTGYRLFEEILGSGSGGSGSGGSGSGDKEWILQKIYEIMRLLDELGHAEASMRVSDLIYEFMK
KGDERLLEEAERLLEEVERGSGSGGSGSGGSGSGMVFTLEDFVGDWEQTAAYNLDQVLEQGGVSSLLQNLAVSVTPIQR
IVRSGENALKIDIHVIIPYEGLSADQMAQIEEVFKVVYPVDDHHFKVILPYGTLVIDGVTPNMLNYFGRPYEGIAVFDGKKITVT
GTLWNGNKIIDERLITPDGSMLFRVTINS-
(Green: 6xHis, Purple: TEVcs, Red: SmBiT, Orange: LCB1, Blue: LgBiT)

**6xHis-TEVcs-SmBiT-15aa-LCB1-10aa-LgBiT**

ATGGCGCATCACCACCACCACCACGAGAATCTTTATTTCCAAGGTTCGGTGACCGGGTACCGTTTGTTCGAGGAGAT
TCTGGGTTCTGGATCAGGAGGGTCTGGAAGCGGCGGCTCAGGTTCCGGAGATAAGGAATGGATTTTGCAAAAGATC
TATGAGATTATGCGTTTACTTGACGAACTTGGACACGCTGAGGCGTCGATGCGTGTGTCGGACCTCATTTATGAGTTC
ATGAAGAAGGGCGACGAGCGCTTGTTGGAAGAGGCCGAGCGTCTCCTCGAAGAGGTCGAGCGCGGTTCAGGAAGC
GGCGGCAGCGGCTCAGGGATGGTATTTACGTTGGAAGATTTCGTGGGAGACTGGGAGCAAACCGCGGCCTATAACT
TAGATCAGGTGTTGGAGCAAGGTGGTGTTTCGAGTTTACTGCAGAACCTTGCCGTGTCCGTCACACCTATTCAGCGT
ATCGTCCGTTCAGGTGAGAATGCACTCAAGATTGACATTCACGTCATCATCCCGTACGAGGGGTTAAGCGCAGACCA
AATGGCTCAAATCGAAGAGGTATTTAAGGTCGTGTATCCAGTCGATGACCACCATTTTAAAGTCATCCTGCCTTATGG
CACATTGGTGATTGACGGGGTTACACCAAATATGTTGAACTATTTTGGACGTCCTTACGAGGGGATCGCCGTGTTCGA
TGGAAAGAAGATCACGGTCACGGGGACGTTATGGAATGGTAACAAAATCATCGATGAGCGTTTAATTACGCCGGACG
GCTCGATGTTATTCCGTGTTACAATCAATAGCTGA
Corresponding to:
MAHHHHHHENLYFQ↓GSVTGYRLFEEILGSGSGGSGSGGSGSGDKEWILQKIYEIMRLLDELGHAEASMRVSDLIYEFMK
KGDERLLEEAERLLEEVERGSGSGGSGSGMVFTLEDFVGDWEQTAAYNLDQVLEQGGVSSLLQNLAVSVTPIQRIVRSG
ENALKIDIHVIIPYEGLSADQMAQIEEVFKVVYPVDDHHFKVILPYGTLVIDGVTPNMLNYFGRPYEGIAVFDGKKITVTGTLW
NGNKIIDERLITPDGSMLFRVTINS-
(Green: 6xHis, Purple: TEVcs, Red: SmBiT, Orange: LCB1, Blue: LgBiT)

**6xHis-TEVcs-SmBiT-15aa-LCB1-5aa-LgBiT**

ATGGCGCATCACCACCACCACCACGAGAATCTTTATTTCCAAGGTTCGGTGACCGGGTACCGTTTGTTCGAGGAGAT
TCTGGGTTCTGGATCAGGAGGGTCTGGAAGCGGCGGCTCAGGTTCCGGAGATAAGGAATGGATTTTGCAAAAGATC
TATGAGATTATGCGTTTACTTGACGAACTTGGACACGCTGAGGCGTCGATGCGTGTGTCGGACCTCATTTATGAGTTC
ATGAAGAAGGGCGACGAGCGCTTGTTGGAAGAGGCCGAGCGTCTCCTCGAAGAGGTCGAGCGCGGTTCAGGAAGC
GGCATGGTATTTACGTTGGAAGATTTCGTGGGAGACTGGGAGCAAACCGCGGCCTATAACTTAGATCAGGTGTTGGA
GCAAGGTGGTGTTTCGAGTTTACTGCAGAACCTTGCCGTGTCCGTCACACCTATTCAGCGTATCGTCCGTTCAGGTG
AGAATGCACTCAAGATTGACATTCACGTCATCATCCCGTACGAGGGGTTAAGCGCAGACCAAATGGCTCAAATCGAA
GAGGTATTTAAGGTCGTGTATCCAGTCGATGACCACCATTTTAAAGTCATCCTGCCTTATGGCACATTGGTGATTGAC
GGGGTTACACCAAATATGTTGAACTATTTTGGACGTCCTTACGAGGGGATCGCCGTGTTCGATGGAAAGAAGATCAC
GGTCACGGGGACGTTATGGAATGGTAACAAAATCATCGATGAGCGTTTAATTACGCCGGACGGCTCGATGTTATTCC
GTGTTACAATCAATAGCTGA
Corresponding to:
MAHHHHHHENLYFQ↓GSVTGYRLFEEILGSGSGGSGSGGSGSGDKEWILQKIYEIMRLLDELGHAEASMRVSDLIYEFMK
KGDERLLEEAERLLEEVERGSGSGMVFTLEDFVGDWEQTAAYNLDQVLEQGGVSSLLQNLAVSVTPIQRIVRSGENALKID
IHVIIPYEGLSADQMAQIEEVFKVVYPVDDHHFKVILPYGTLVIDGVTPNMLNYFGRPYEGIAVFDGKKITVTGTLWNGNKIID
ERLITPDGSMLFRVTINS-
(Green: 6xHis, Purple: TEVcs, Red: SmBiT, Orange: LCB1, Blue: LgBiT)

**6xHis-TEVcs-SmBiT-15aa-LCB1-0aa-LgBiT**

ATGGCGCATCACCACCACCACCACGAGAATCTTTATTTCCAAGGTTCGGTGACCGGGTACCGTTTGTTCGAGGAGAT
TCTGGGTTCTGGATCAGGAGGGTCTGGAAGCGGCGGCTCAGGTTCCGGAGATAAGGAATGGATTTTGCAAAAGATC
TATGAGATTATGCGTTTACTTGACGAACTTGGACACGCTGAGGCGTCGATGCGTGTGTCGGACCTCATTTATGAGTTC
ATGAAGAAGGGCGACGAGCGCTTGTTGGAAGAGGCCGAGCGTCTCCTCGAAGAGGTCGAGCGCATGGTATTTACGT
TGGAAGATTTCGTGGGAGACTGGGAGCAAACCGCGGCCTATAACTTAGATCAGGTGTTGGAGCAAGGTGGTGTTTC
GAGTTTACTGCAGAACCTTGCCGTGTCCGTCACACCTATTCAGCGTATCGTCCGTTCAGGTGAGAATGCACTCAAGAT
TGACATTCACGTCATCATCCCGTACGAGGGGTTAAGCGCAGACCAAATGGCTCAAATCGAAGAGGTATTTAAGGTCG
TGTATCCAGTCGATGACCACCATTTTAAAGTCATCCTGCCTTATGGCACATTGGTGATTGACGGGGTTACACCAAATA
TGTTGAACTATTTTGGACGTCCTTACGAGGGGATCGCCGTGTTCGATGGAAAGAAGATCACGGTCACGGGGACGTTA
TGGAATGGTAACAAAATCATCGATGAGCGTTTAATTACGCCGGACGGCTCGATGTTATTCCGTGTTACAATCAATAGC
TGA

Corresponding to:

MAHHHHHHENLYFQ↓GSVTGYRLFEEILGSGSGGSGSGGSGSGDKEWILQKIYEIMRLLDELGHAEASMRVSDLIYEFMK
KGDERLLEEAERLLEEVERMVFTLEDFVGDWEQTAAYNLDQVLEQGGVSSLLQNLAVSVTPIQRIVRSGENALKIDIHVIIPY
EGLSADQMAQIEEVFKVVYPVDDHHFKVILPYGTLVIDGVTPNMLNYFGRPYEGIAVFDGKKITVTGTLWNGNKIIDERLITP
DGSMLFRVTINS-

(Green: 6xHis, Purple: TEVcs, Red: SmBiT, Orange: LCB1, Blue: LgBiT)

**<u>6xHis-TEVcs-SmBiT-10aa-LCB1-15aa-LgBiT</u>**

ATGGCGCATCACCACCACCACCACGAGAATCTTTATTTCCAAGGTTCGGTGACCGGGTACCGTTTGTTCGAGGAGAT
TCTGGGGTCTGGAAGCGGCGGCTCAGGTTCCGGAGATAAGGAATGGATTTTGCAAAAGATCTATGAGATTATGCGTT
TACTTGACGAACTTGGACACGCTGAGGCGTCGATGCGTGTGTCGGACCTCATTTATGAGTTCATGAAGAAGGGCGAC
GAGCGCTTGTTGGAAGAGGCCGAGCGTCTCCTCGAAGAGGTCGAGCGCGGTTCTGGATCAGGAGGTTCAGGAAGC
GGCGGCAGCGGCTCAGGGATGGTATTTACGTTGGAAGATTTCGTGGGAGACTGGGAGCAAACCGCGGCCTATAACT
TAGATCAGGTGTTGGAGCAAGGTGGTGTTTCGAGTTTACTGCAGAACCTTGCCGTGTCCGTCACACCTATTCAGCGT
ATCGTCCGTTCAGGTGAGAATGCACTCAAGATTGACATTCACGTCATCATCCCGTACGAGGGGTTAAGCGCAGACCA
AATGGCTCAAATCGAAGAGGTATTTAAGGTCGTGTATCCAGTCGATGACCACCATTTTAAAGTCATCCTGCCTTATGG
CACATTGGTGATTGACGGGGTTACACCAAATATGTTGAACTATTTTGGACGTCCTTACGAGGGGATCGCCGTGTTCGA
TGGAAAGAAGATCACGGTCACGGGGACGTTATGGAATGGTAACAAAATCATCGATGAGCGTTTAATTACGCCGGACG
GCTCGATGTTATTCCGTGTTACAATCAATAGCTGA

Corresponding to:

MAHHHHHHENLYFQ↓GSVTGYRLFEEILGSGSGGSGSGDKEWILQKIYEIMRLLDELGHAEASMRVSDLIYEFMKKGDERL
LEEAERLLEEVERGSGSGGSGSGGSGSGMVFTLEDFVGDWEQTAAYNLDQVLEQGGVSSLLQNLAVSVTPIQRIVRSGE
NALKIDIHVIIPYEGLSADQMAQIEEVFKVVYPVDDHHFKVILPYGTLVIDGVTPNMLNYFGRPYEGIAVFDGKKITVTGTLWN
GNKIIDERLITPDGSMLFRVTINS-

(Green: 6xHis, Purple: TEVcs, Red: SmBiT, Orange: LCB1, Blue: LgBiT)

**<u>6xHis-TEVcs-SmBiT-10aa-LCB1-10aa-LgBiT</u>**

ATGGCGCATCACCACCACCACCACGAGAATCTTTATTTCCAAGGTTCGGTGACCGGGTACCGTTTGTTCGAGGAGAT
TCTGGGGTCTGGAAGCGGCGGCTCAGGTTCCGGAGATAAGGAATGGATTTTGCAAAAGATCTATGAGATTATGCGTT
TACTTGACGAACTTGGACACGCTGAGGCGTCGATGCGTGTGTCGGACCTCATTTATGAGTTCATGAAGAAGGGCGAC
GAGCGCTTGTTGGAAGAGGCCGAGCGTCTCCTCGAAGAGGTCGAGCGCGGTTCAGGAAGCGGCGGCAGCGGCTCA
GGGATGGTATTTACGTTGGAAGATTTCGTGGGAGACTGGGAGCAAACCGCGGCCTATAACTTAGATCAGGTGTTGGA
GCAAGGTGGTGTTTCGAGTTTACTGCAGAACCTTGCCGTGTCCGTCACACCTATTCAGCGTATCGTCCGTTCAGGTG
AGAATGCACTCAAGATTGACATTCACGTCATCATCCCGTACGAGGGGTTAAGCGCAGACCAAATGGCTCAAATCGAA
GAGGTATTTAAGGTCGTGTATCCAGTCGATGACCACCATTTTAAAGTCATCCTGCCTTATGGCACATTGGTGATTGAC
GGGGTTACACCAAATATGTTGAACTATTTTGGACGTCCTTACGAGGGGATCGCCGTGTTCGATGGAAAGAAGATCAC
GGTCACGGGGACGTTATGGAATGGTAACAAAATCATCGATGAGCGTTTAATTACGCCGGACGGCTCGATGTTATTCC
GTGTTACAATCAATAGCTGA

Corresponding to:

MAHHHHHHENLYFQ↓GSVTGYRLFEEILGSGSGGSGSGDKEWILQKIYEIMRLLDELGHAEASMRVSDLIYEFMKKGDERL
LEEAERLLEEVERGSGSGGSGSGMVFTLEDFVGDWEQTAAYNLDQVLEQGGVSSLLQNLAVSVTPIQRIVRSGENALKIDI
HVIIPYEGLSADQMAQIEEVFKVVYPVDDHHFKVILPYGTLVIDGVTPNMLNYFGRPYEGIAVFDGKKITVTGTLWNGNKIIDE
RLITPDGSMLFRVTINS-

(Green: 6xHis, Purple: TEVcs, Red: SmBiT, Orange: LCB1, Blue: LgBiT)

**<u>6xHis-TEVcs-SmBiT-10aa-LCB1-5aa-LgBiT</u>**

ATGGCGCATCACCACCACCACCACGAGAATCTTTATTTCCAAGGTTCGGTGACCGGGTACCGTTTGTTCGAGGAGAT
TCTGGGGTCTGGAAGCGGCGGCTCAGGTTCCGGAGATAAGGAATGGATTTTGCAAAAGATCTATGAGATTATGCGTT
TACTTGACGAACTTGGACACGCTGAGGCGTCGATGCGTGTGTCGGACCTCATTTATGAGTTCATGAAGAAGGGCGAC
GAGCGCTTGTTGGAAGAGGCCGAGCGTCTCCTCGAAGAGGTCGAGCGCGGTTCAGGAAGCGGCATGGTATTTACGT

TGGAAGATTTCGTGGGAGACTGGGAGCAAACCGCGGCCTATAACTTAGATCAGGTGTTGGAGCAAGGTGGTGTTTC
GAGTTTACTGCAGAACCTTGCCGTGTCCGTCACACCTATTCAGCGTATCGTCCGTTCAGGTGAGAATGCACTCAAGAT
TGACATTCACGTCATCATCCCGTACGAGGGGTTAAGCGCAGACCAAATGGCTCAAATCGAAGAGGTATTTAAGGTCG
TGTATCCAGTCGATGACCACCATTTTAAAGTCATCCTGCCTTATGGCACATTGGTGATTGACGGGGTTACACCAAATA
TGTTGAACTATTTTGGACGTCCTTACGAGGGGATCGCCGTGTTCGATGGAAAGAAGATCACGGTCACGGGGACGTTA
TGGAATGGTAACAAAATCATCGATGAGCGTTTAATTACGCCGGACGGCTCGATGTTATTCCGTGTTACAATCAATAGC
TGA
Corresponding to:
MAHHHHHHENLYFQ↓GSVTGYRLFEEILGSGSGGSGSGDKEWILQKIYEIMRLLDELGHAEASMRVSDLIYEFMKKGDERL
LEEAERLLEEVERGSGSGMVFTLEDFVGDWEQTAAYNLDQVLEQGGVSSLLQNLAVSVTPIQRIVRSGENALKIDIHVIIPY
EGLSADQMAQIEEVFKVVYPVDDHHFKVILPYGTLVIDGVTPNMLNYFGRPYEGIAVFDGKKITVTGTLWNGNKIIDERLITP
DGSMLFRVTINS-
(Green: 6xHis, Purple: TEVcs, Red: SmBiT, Orange: LCB1, Blue: LgBiT)

**6xHis-TEVcs-SmBiT-10aa-LCB1-0aa-LgBiT**
ATGGCGCATCACCACCACCACCACGAGAATCTTTATTTCCAAGGTTCGGTGACCGGGTACCGTTTGTTCGAGGAGAT
TCTGGGGTCTGGAAGCGGCGGCTCAGGTTCCGGAGATAAGGAATGGATTTTGCAAAAGATCTATGAGATTATGCGTT
TACTTGACGAACTTGGACACGCTGAGGCGTCGATGCGTGTGTCGGACCTCATTTATGAGTTCATGAAGAAGGGCGAC
GAGCGCTTGTTGGAAGAGGCCGAGCGTCTCCTCGAAGAGGTCGAGCGCATGGTATTTACGTTGGAAGATTTCGTGG
GAGACTGGGAGCAAACCGCGGCCTATAACTTAGATCAGGTGTTGGAGCAAGGTGGTGTTTCGAGTTTACTGCAGAAC
CTTGCCGTGTCCGTCACACCTATTCAGCGTATCGTCCGTTCAGGTGAGAATGCACTCAAGATTGACATTCACGTCATC
ATCCCGTACGAGGGGTTAAGCGCAGACCAAATGGCTCAAATCGAAGAGGTATTTAAGGTCGTGTATCCAGTCGATGA
CCACCATTTTAAAGTCATCCTGCCTTATGGCACATTGGTGATTGACGGGGTTACACCAAATATGTTGAACTATTTTGGA
CGTCCTTACGAGGGGATCGCCGTGTTCGATGGAAAGAAGATCACGGTCACGGGGACGTTATGGAATGGTAACAAAA
TCATCGATGAGCGTTTAATTACGCCGGACGGCTCGATGTTATTCCGTGTTACAATCAATAGCTGA
Corresponding to:
MAHHHHHHENLYFQ↓GSVTGYRLFEEILGSGSGGSGSGDKEWILQKIYEIMRLLDELGHAEASMRVSDLIYEFMKKGDERL
LEEAERLLEEVERMVFTLEDFVGDWEQTAAYNLDQVLEQGGVSSLLQNLAVSVTPIQRIVRSGENALKIDIHVIIPYEGLSAD
QMAQIEEVFKVVYPVDDHHFKVILPYGTLVIDGVTPNMLNYFGRPYEGIAVFDGKKITVTGTLWNGNKIIDERLITPDGSMLF
RVTINS-
(Green: 6xHis, Purple: TEVcs, Red: SmBiT, Orange: LCB1, Blue: LgBiT)

**6xHis-TEVcs-SmBiT-5aa-LCB1-15aa-LgBiT**
ATGGCGCATCACCACCACCACCACGAGAATCTTTATTTCCAAGGTTCGGTGACCGGGTACCGTTTGTTCGAGGAGAT
TCTGGGGTCTGGAAGCGGCGATAAGGAATGGATTTTGCAAAAGATCTATGAGATTATGCGTTTACTTGACGAACTTGG
ACACGCTGAGGCGTCGATGCGTGTGTCGGACCTCATTTATGAGTTCATGAAGAAGGGCGACGAGCGCTTGTTGGAA
GAGGCCGAGCGTCTCCTCGAAGAGGTCGAGCGCGGTTCTGGATCAGGAGGTTCAGGAAGCGGCGGCAGCGGCTCA
GGGATGGTATTTACGTTGGAAGATTTCGTGGGAGACTGGGAGCAAACCGCGGCCTATAACTTAGATCAGGTGTTGGA
GCAAGGTGGTGTTTCGAGTTTACTGCAGAACCTTGCCGTGTCCGTCACACCTATTCAGCGTATCGTCCGTTCAGGTG
AGAATGCACTCAAGATTGACATTCACGTCATCATCCCGTACGAGGGGTTAAGCGCAGACCAAATGGCTCAAATCGAA
GAGGTATTTAAGGTCGTGTATCCAGTCGATGACCACCATTTTAAAGTCATCCTGCCTTATGGCACATTGGTGATTGAC
GGGGTTACACCAAATATGTTGAACTATTTTGGACGTCCTTACGAGGGGATCGCCGTGTTCGATGGAAAGAAGATCAC
GGTCACGGGGACGTTATGGAATGGTAACAAAATCATCGATGAGCGTTTAATTACGCCGGACGGCTCGATGTTATTCC
GTGTTACAATCAATAGCTGA
Corresponding to:
MAHHHHHHENLYFQ↓GSVTGYRLFEEILGSGSGDKEWILQKIYEIMRLLDELGHAEASMRVSDLIYEFMKKGDERLLEEAE
RLLEEVERGSGSGGSGSGGSGSGMVFTLEDFVGDWEQTAAYNLDQVLEQGGVSSLLQNLAVSVTPIQRIVRSGENALKID
IHVIIPYEGLSADQMAQIEEVFKVVYPVDDHHFKVILPYGTLVIDGVTPNMLNYFGRPYEGIAVFDGKKITVTGTLWNGNKIID
ERLITPDGSMLFRVTINS-
(Green: 6xHis, Purple: TEVcs, Red: SmBiT, Orange: LCB1, Blue: LgBiT)

**6xHis-TEVcs-SmBiT-5aa-LCB1-10aa-LgBiT**
ATGGCGCATCACCACCACCACCACGAGAATCTTTATTTCCAAGGTTCGGTGACCGGGTACCGTTTGTTCGAGGAGAT
TCTGGGGTCTGGAAGCGGCGATAAGGAATGGATTTTGCAAAAGATCTATGAGATTATGCGTTTACTTGACGAACTTGG
ACACGCTGAGGCGTCGATGCGTGTGTCGGACCTCATTTATGAGTTCATGAAGAAGGGCGACGAGCGCTTGTTGGAA
GAGGCCGAGCGTCTCCTCGAAGAGGTCGAGCGCGGTTCAGGAAGCGGCGGCAGCGGCTCAGGGATGGTATTTACG
TTGGAAGATTTCGTGGGAGACTGGGAGCAAACCGCGGCCTATAACTTAGATCAGGTGTTGGAGCAAGGTGGTGTTTC
GAGTTTACTGCAGAACCTTGCCGTGTCCGTCACACCTATTCAGCGTATCGTCCGTTCAGGTGAGAATGCACTCAAGAT
TGACATTCACGTCATCATCCCGTACGAGGGGTTAAGCGCAGACCAAATGGCTCAAATCGAAGAGGTATTTAAGGTCG
TGTATCCAGTCGATGACCACCATTTTAAAGTCATCCTGCCTTATGGCACATTGGTGATTGACGGGGTTACACCAAATA
TGTTGAACTATTTTGGACGTCCTTACGAGGGGATCGCCGTGTTCGATGGAAAGAAGATCACGGTCACGGGGACGTTA

TGGAATGGTAACAAAATCATCGATGAGCGTTTAATTACGCCGGACGGCTCGATGTTATTCCGTGTTACAATCAATAGC
TGA
Corresponding to:
MAHHHHHHENLYFQ↓GSVTGYRLFEEILGSGSGDKEWILQKIYEIMRLLDELGHAEASMRVSDLIYEFMKKGDERLLEEAERLLEEVERGSGSGGSGSGMVFTLEDFVGDWEQTAAYNLDQVLEQGGVSSLLQNLAVSVTPIQRIVRSGENALKIDIHVIIPYEGLSADQMAQIEEVFKVVYPVDDHHFKVILPYGTLVIDGVTPNMLNYFGRPYEGIAVFDGKKITVTGTLWNGNKIIDERLITPDGSMLFRVTINS-
(Green: 6xHis, Purple: TEVcs, Red: SmBiT, Orange: LCB1, Blue: LgBiT)

**6xHis-TEVcs-SmBiT-5aa-LCB1-5aa-LgBiT**

ATGGCGCATCACCACCACCACCACGAGAATCTTTATTTCCAAGGTTCGGTGACCGGGTACCGTTTGTTCGAGGAGATTCTGGGGTCTGGAAGCGGCGATAAGGAATGGATTTTGCAAAAGATCTATGAGATTATGCGTTTACTTGACGAACTTGGACACGCTGAGGCGTCGATGCGTGTGTCGGACCTCATTTATGAGTTCATGAAGAAGGGCGACGAGCGCTTGTTGGAAGAGGCCGAGCGTCTCCTCGAAGAGGTCGAGCGCGGTTCAGGAAGCGGCATGGTATTTACGTTGGAAGATTTCGTGGGAGACTGGGAGCAAACCGCGGCCTATAACTTAGATCAGGTGTTGGAGCAAGGTGGTGTTTCGAGTTTACTGCAGAACCTTGCCGTGTCCGTCACACCTATTCAGCGTATCGTCCGTTCAGGTGAGAATGCACTCAAGATTGACATTCACGTCATCATCCCGTACGAGGGGTTAAGCGCAGACCAAATGGCTCAAATCGAAGAGGTATTTAAGGTCGTGTATCCAGTCGATGACCACCATTTTAAAGTCATCCTGCCTTATGGCACATTGGTGATTGACGGGGTTACACCAAATATGTTGAACTATTTTGGACGTCCTTACGAGGGGATCGCCGTGTTCGATGGAAAGAAGATCACGGTCACGGGGACGTTATGGAATGGTAACAAAATCATCGATGAGCGTTTAATTACGCCGGACGGCTCGATGTTATTCCGTGTTACAATCAATAGCTGA
Corresponding to:
MAHHHHHHENLYFQ↓GSVTGYRLFEEILGSGSGDKEWILQKIYEIMRLLDELGHAEASMRVSDLIYEFMKKGDERLLEEAERLLEEVERGSGSGMVFTLEDFVGDWEQTAAYNLDQVLEQGGVSSLLQNLAVSVTPIQRIVRSGENALKIDIHVIIPYEGLSADQMAQIEEVFKVVYPVDDHHFKVILPYGTLVIDGVTPNMLNYFGRPYEGIAVFDGKKITVTGTLWNGNKIIDERLITPDGSMLFRVTINS-
(Green: 6xHis, Purple: TEVcs, Red: SmBiT, Orange: LCB1, Blue: LgBiT)

**6xHis-TEVcs-SmBiT-5aa-LCB1-0aa-LgBiT**

ATGGCGCATCACCACCACCACCACGAGAATCTTTATTTCCAAGGTTCGGTGACCGGGTACCGTTTGTTCGAGGAGATTCTGGGGTCTGGAAGCGGCGATAAGGAATGGATTTTGCAAAAGATCTATGAGATTATGCGTTTACTTGACGAACTTGGACACGCTGAGGCGTCGATGCGTGTGTCGGACCTCATTTATGAGTTCATGAAGAAGGGCGACGAGCGCTTGTTGGAAGAGGCCGAGCGTCTCCTCGAAGAGGTCGAGCGCATGGTATTTACGTTGGAAGATTTCGTGGGAGACTGGGAGCAAACCGCGGCCTATAACTTAGATCAGGTGTTGGAGCAAGGTGGTGTTTCGAGTTTACTGCAGAACCTTGCCGTGTCCGTCACACCTATTCAGCGTATCGTCCGTTCAGGTGAGAATGCACTCAAGATTGACATTCACGTCATCATCCCGTACGAGGGGTTAAGCGCAGACCAAATGGCTCAAATCGAAGAGGTATTTAAGGTCGTGTATCCAGTCGATGACCACCATTTTAAAGTCATCCTGCCTTATGGCACATTGGTGATTGACGGGGTTACACCAAATATGTTGAACTATTTTGGACGTCCTTACGAGGGGATCGCCGTGTTCGATGGAAAGAAGATCACGGTCACGGGGACGTTATGGAATGGTAACAAAATCATCGATGAGCGTTTAATTACGCCGGACGGCTCGATGTTATTCCGTGTTACAATCAATAGCTGA
Corresponding to:
MAHHHHHHENLYFQ↓GSVTGYRLFEEILGSGSGDKEWILQKIYEIMRLLDELGHAEASMRVSDLIYEFMKKGDERLLEEAERLLEEVERMVFTLEDFVGDWEQTAAYNLDQVLEQGGVSSLLQNLAVSVTPIQRIVRSGENALKIDIHVIIPYEGLSADQMAQIEEVFKVVYPVDDHHFKVILPYGTLVIDGVTPNMLNYFGRPYEGIAVFDGKKITVTGTLWNGNKIIDERLITPDGSMLFRVTINS
-
(Green: 6xHis, Purple: TEVcs, Red: SmBiT, Orange: LCB1, Blue: LgBiT)

**6xHis-TEVcs-SmBiT-0aa-LCB1-15aa-LgBiT**

ATGGCGCATCACCACCACCACCACGAGAATCTTTATTTCCAAGGTTCGGTGACCGGGTACCGTTTGTTCGAGGAGATTCTGGATAAGGAATGGATTTTGCAAAAGATCTATGAGATTATGCGTTTACTTGACGAACTTGGACACGCTGAGGCGTCGATGCGTGTGTCGGACCTCATTTATGAGTTCATGAAGAAGGGCGACGAGCGCTTGTTGGAAGAGGCCGAGCGTCTCCTCGAAGAGGTCGAGCGCGGTTCTGGATCAGGAGGTTCAGGAAGCGGCGGCAGCGGCTCAGGGATGGTATTTACGTTGGAAGATTTCGTGGGAGACTGGGAGCAAACCGCGGCCTATAACTTAGATCAGGTGTTGGAGCAAGGTGGTGTTTCGAGTTTACTGCAGAACCTTGCCGTGTCCGTCACACCTATTCAGCGTATCGTCCGTTCAGGTGAGAATGCACTCAAGATTGACATTCACGTCATCATCCCGTACGAGGGGTTAAGCGCAGACCAAATGGCTCAAATCGAAGAGGTATTTAAGGTCGTGTATCCAGTCGATGACCACCATTTTAAAGTCATCCTGCCTTATGGCACATTGGTGATTGACGGGGTTACACCAAATATGTTGAACTATTTTGGACGTCCTTACGAGGGGATCGCCGTGTTCGATGGAAAGAAGATCACGGTCACGGGGACGTTATGGAATGGTAACAAAATCATCGATGAGCGTTTAATTACGCCGGACGGCTCGATGTTATTCCGTGTTACAATCAATAGC
TGA
Corresponding to:
MAHHHHHHENLYFQ↓GSVTGYRLFEEILDKEWILQKIYEIMRLLDELGHAEASMRVSDLIYEFMKKGDERLLEEAERLLEEVERGSGSGGSGSGGSGSGMVFTLEDFVGDWEQTAAYNLDQVLEQGGVSSLLQNLAVSVTPIQRIVRSGENALKIDIHVIIPY

EGLSADQMAQIEEVFKVVYPVDDHHFKVILPYGTLVIDGVTPNMLNYFGRPYEGIAVFDGKKITVTGTLWNGNKIIDERLITP
DGSMLFRVTINS-
(Green: 6xHis, Purple: TEVcs, Red: SmBiT, Orange: LCB1, Blue: LgBiT)

**6xHis-TEVcs-SmBiT-0aa-LCB1-10aa-LgBiT**
ATGGCGCATCACCACCACCACCACGAGAATCTTTATTTCCAAGGTTCGGTGACCGGGTACCGTTTGTTCGAGGAGAT
TCTGGATAAGGAATGGATTTTGCAAAAGATCTATGAGATTATGCGTTTACTTGACGAACTTGGACACGCTGAGGCGTC
GATGCGTGTGTCGGACCTCATTTATGAGTTCATGAAGAAGGGCGACGAGCGCTTGTTGGAAGAGGCCGAGCGTCTC
CTCGAAGAGGTCGAGCGCGGTTCAGGAAGCGGCGGCAGCGGCTCAGGGATGGTATTTACGTTGGAAGATTTCGTG
GGAGACTGGGAGCAAACCGCGGCCTATAACTTAGATCAGGTGTTGGAGCAAGGTGGTGTTTCGAGTTTACTGCAGAA
CCTTGCCGTGTCCGTCACACCTATTCAGCGTATCGTCCGTTCAGGTGAGAATGCACTCAAGATTGACATTCACGTCAT
CATCCCGTACGAGGGGTTAAGCGCAGACCAAATGGCTCAAATCGAAGAGGTATTTAAGGTCGTGTATCCAGTCGATG
ACCACCATTTTAAAGTCATCCTGCCTTATGGCACATTGGTGATTGACGGGGTTACACCAAATATGTTGAACTATTTTGG
ACGTCCTTACGAGGGGATCGCCGTGTTCGATGGAAAGAAGATCACGGTCACGGGGACGTTATGGAATGGTAACAAA
ATCATCGATGAGCGTTTAATTACGCCGGACGGCTCGATGTTATTCCGTGTTACAATCAATAGCTGA
Corresponding to:
MAHHHHHHENLYFQ↓GSVTGYRLFEEILDKEWILQKIYEIMRLLDELGHAEASMRVSDLIYEFMKKGDERLLEEAERLLEEV
ERGSGSGGSGSGMVFTLEDFVGDWEQTAAYNLDQVLEQGGVSSLLQNLAVSVTPIQRIVRSGENALKIDIHVIIPYEGLSA
DQMAQIEEVFKVVYPVDDHHFKVILPYGTLVIDGVTPNMLNYFGRPYEGIAVFDGKKITVTGTLWNGNKIIDERLITPDGSML
FRVTINS-
(Green: 6xHis, Purple: TEVcs, Red: SmBiT, Orange: LCB1, Blue: LgBiT)

**6xHis-TEVcs-SmBiT-0aa-LCB1-5aa-LgBiT**
ATGGCGCATCACCACCACCACCACGAGAATCTTTATTTCCAAGGTTCGGTGACCGGGTACCGTTTGTTCGAGGAGAT
TCTGGATAAGGAATGGATTTTGCAAAAGATCTATGAGATTATGCGTTTACTTGACGAACTTGGACACGCTGAGGCGTC
GATGCGTGTGTCGGACCTCATTTATGAGTTCATGAAGAAGGGCGACGAGCGCTTGTTGGAAGAGGCCGAGCGTCTC
CTCGAAGAGGTCGAGCGCGGTTCAGGAAGCGGCATGGTATTTACGTTGGAAGATTTCGTGGGAGACTGGGAGCAAA
CCGCGGCCTATAACTTAGATCAGGTGTTGGAGCAAGGTGGTGTTTCGAGTTTACTGCAGAACCTTGCCGTGTCCGTC
ACACCTATTCAGCGTATCGTCCGTTCAGGTGAGAATGCACTCAAGATTGACATTCACGTCATCATCCCGTACGAGGG
GTTAAGCGCAGACCAAATGGCTCAAATCGAAGAGGTATTTAAGGTCGTGTATCCAGTCGATGACCACCATTTTAAAGT
CATCCTGCCTTATGGCACATTGGTGATTGACGGGGTTACACCAAATATGTTGAACTATTTTGGACGTCCTTACGAGGG
GATCGCCGTGTTCGATGGAAAGAAGATCACGGTCACGGGGACGTTATGGAATGGTAACAAAATCATCGATGAGCGTT
TAATTACGCCGGACGGCTCGATGTTATTCCGTGTTACAATCAATAGCTGA
Corresponding to:
MAHHHHHHENLYFQ↓GSVTGYRLFEEILDKEWILQKIYEIMRLLDELGHAEASMRVSDLIYEFMKKGDERLLEEAERLLEEV
ERGSGSGMVFTLEDFVGDWEQTAAYNLDQVLEQGGVSSLLQNLAVSVTPIQRIVRSGENALKIDIHVIIPYEGLSADQMAQI
EEVFKVVYPVDDHHFKVILPYGTLVIDGVTPNMLNYFGRPYEGIAVFDGKKITVTGTLWNGNKIIDERLITPDGSMLFRVTINS
-
(Green: 6xHis, Purple: TEVcs, Red: SmBiT, Orange: LCB1, Blue: LgBiT)

**6xHis-TEVcs-SmBiT-0aa-LCB1-0aa-LgBiT**
ATGGCGCATCACCACCACCACCACGAGAATCTTTATTTCCAAGGTTCGGTGACCGGGTACCGTTTGTTCGAGGAGAT
TCTGGATAAGGAATGGATTTTGCAAAAGATCTATGAGATTATGCGTTTACTTGACGAACTTGGACACGCTGAGGCGTC
GATGCGTGTGTCGGACCTCATTTATGAGTTCATGAAGAAGGGCGACGAGCGCTTGTTGGAAGAGGCCGAGCGTCTC
CTCGAAGAGGTCGAGCGCATGGTATTTACGTTGGAAGATTTCGTGGGAGACTGGGAGCAAACCGCGGCCTATAACTT
AGATCAGGTGTTGGAGCAAGGTGGTGTTTCGAGTTTACTGCAGAACCTTGCCGTGTCCGTCACACCTATTCAGCGTA
TCGTCCGTTCAGGTGAGAATGCACTCAAGATTGACATTCACGTCATCATCCCGTACGAGGGGTTAAGCGCAGACCAA
ATGGCTCAAATCGAAGAGGTATTTAAGGTCGTGTATCCAGTCGATGACCACCATTTTAAAGTCATCCTGCCTTATGGC
ACATTGGTGATTGACGGGGTTACACCAAATATGTTGAACTATTTTGGACGTCCTTACGAGGGGATCGCCGTGTTCGAT
GGAAAGAAGATCACGGTCACGGGGACGTTATGGAATGGTAACAAAATCATCGATGAGCGTTTAATTACGCCGGACGG
CTCGATGTTATTCCGTGTTACAATCAATAGCTGA
Corresponding to:
MAHHHHHHENLYFQ↓GSVTGYRLFEEILDKEWILQKIYEIMRLLDELGHAEASMRVSDLIYEFMKKGDERLLEEAERLLEEV
ERMVFTLEDFVGDWEQTAAYNLDQVLEQGGVSSLLQNLAVSVTPIQRIVRSGENALKIDIHVIIPYEGLSADQMAQIEEVFK
VVYPVDDHHFKVILPYGTLVIDGVTPNMLNYFGRPYEGIAVFDGKKITVTGTLWNGNKIIDERLITPDGSMLFRVTINS-
(Green: 6xHis, Purple: TEVcs, Red: SmBiT, Orange: LCB1, Blue: LgBiT)

**6xHis-TEVcs-SmBiT-4aa-LCB1-0aa-LgBiT**
ATGGCGCATCACCACCACCACCACGAGAATCTTTATTTCCAAGGTTCGGTGACCGGGTACCGTTTGTTCGAGGAGAT
TCTGGGTTCTGGATCAGATAAGGAATGGATTTTGCAAAAGATCTATGAGATTATGCGTTTACTTGACGAACTTGGACA
CGCTGAGGCGTCGATGCGTGTGTCGGACCTCATTTATGAGTTCATGAAGAAGGGCGACGAGCGCTTGTTGGAAGAG

GCCGAGCGTCTCCTCGAAGAGGTCGAGCGCATGGTATTTACGTTGGAAGATTTCGTGGGAGACTGGGAGCAAACCG
CGGCCTATAACTTAGATCAGGTGTTGGAGCAAGGTGGTGTTTCGAGTTTACTGCAGAACCTTGCCGTGTCCGTCACA
CCTATTCAGCGTATCGTCCGTTCAGGTGAGAATGCACTCAAGATTGACATTCACGTCATCATCCCGTACGAGGGGTTA
AGCGCAGACCAAATGGCTCAAATCGAAGAGGTATTTAAGGTCGTGTATCCAGTCGATGACCACCATTTTAAAGTCATC
CTGCCTTATGGCACATTGGTGATTGACGGGGTTACACCAAATATGTTGAACTATTTTGGACGTCCTTACGAGGGGATC
GCCGTGTTCGATGGAAAGAAGATCACGGTCACGGGGACGTTATGGAATGGTAACAAAATCATCGATGAGCGTTTAAT
TACGCCGGACGGCTCGATGTTATTCCGTGTTACAATCAATAGCTGA

Corresponding to:

MAHHHHHHENLYFQ↓GSVTGYRLFEEILGSGSDKEWILQKIYEIMRLLDELGHAEASMRVSDLIYEFMKKGDERLLEEAERL
LEEVERMVFTLEDFVGDWEQTAAYNLDQVLEQGGVSSLLQNLAVSVTPIQRIVRSGENALKIDIHVIIPYEGLSADQMAQIEE
VFKVVYPVDDHHFKVILPYGTLVIDGVTPNMLNYFGRPYEGIAVFDGKKITVTGTLWNGNKIIDERLITPDGSMLFRVTINS-
(Green: 6xHis, Purple: TEVcs, Red: SmBiT, Orange: LCB1, Blue: LgBiT)

**<u>6xHis-TEVcs-SmBiT-3aa-LCB1-0aa-LgBiT</u>**

ATGGCGCATCACCACCACCACCACGAGAATCTTTATTTCCAAGGTTCGGTGACCGGGTACCGTTTGTTCGAGGAGAT
TCTGGGTTCTGGAGATAAGGAATGGATTTTGCAAAAGATCTATGAGATTATGCGTTTACTTGACGAACTTGGACACGC
TGAGGCGTCGATGCGTGTGTCGGACCTCATTTATGAGTTCATGAAGAAGGGCGACGAGCGCTTGTTGGAAGAGGCC
GAGCGTCTCCTCGAAGAGGTCGAGCGCATGGTATTTACGTTGGAAGATTTCGTGGGAGACTGGGAGCAAACCGCGG
CCTATAACTTAGATCAGGTGTTGGAGCAAGGTGGTGTTTCGAGTTTACTGCAGAACCTTGCCGTGTCCGTCACACCTA
TTCAGCGTATCGTCCGTTCAGGTGAGAATGCACTCAAGATTGACATTCACGTCATCATCCCGTACGAGGGGTTAAGC
GCAGACCAAATGGCTCAAATCGAAGAGGTATTTAAGGTCGTGTATCCAGTCGATGACCACCATTTTAAAGTCATCCTG
CCTTATGGCACATTGGTGATTGACGGGGTTACACCAAATATGTTGAACTATTTTGGACGTCCTTACGAGGGGATCGCC
GTGTTCGATGGAAAGAAGATCACGGTCACGGGGACGTTATGGAATGGTAACAAAATCATCGATGAGCGTTTAATTAC
GCCGGACGGCTCGATGTTATTCCGTGTTACAATCAATAGCTGA

Corresponding to:

MAHHHHHHENLYFQ↓GSVTGYRLFEEILGSGDKEWILQKIYEIMRLLDELGHAEASMRVSDLIYEFMKKGDERLLEEAERLL
EEVERMVFTLEDFVGDWEQTAAYNLDQVLEQGGVSSLLQNLAVSVTPIQRIVRSGENALKIDIHVIIPYEGLSADQMAQIEE
VFKVVYPVDDHHFKVILPYGTLVIDGVTPNMLNYFGRPYEGIAVFDGKKITVTGTLWNGNKIIDERLITPDGSMLFRVTINS-
(Green: 6xHis, Purple: TEVcs, Red: SmBiT, Orange: LCB1, Blue: LgBiT)

**<u>6xHis-TEVcs-SmBiT-2aa-LCB1-0aa-LgBiT</u>**

ATGGCGCATCACCACCACCACCACGAGAATCTTTATTTCCAAGGTTCGGTGACCGGGTACCGTTTGTTCGAGGAGAT
TCTGGGTTCTGATAAGGAATGGATTTTGCAAAAGATCTATGAGATTATGCGTTTACTTGACGAACTTGGACACGCTGA
GGCGTCGATGCGTGTGTCGGACCTCATTTATGAGTTCATGAAGAAGGGCGACGAGCGCTTGTTGGAAGAGGCCGAG
CGTCTCCTCGAAGAGGTCGAGCGCATGGTATTTACGTTGGAAGATTTCGTGGGAGACTGGGAGCAAACCGCGGCCT
ATAACTTAGATCAGGTGTTGGAGCAAGGTGGTGTTTCGAGTTTACTGCAGAACCTTGCCGTGTCCGTCACACCTATTC
AGCGTATCGTCCGTTCAGGTGAGAATGCACTCAAGATTGACATTCACGTCATCATCCCGTACGAGGGGTTAAGCGCA
GACCAAATGGCTCAAATCGAAGAGGTATTTAAGGTCGTGTATCCAGTCGATGACCACCATTTTAAAGTCATCCTGCCT
TATGGCACATTGGTGATTGACGGGGTTACACCAAATATGTTGAACTATTTTGGACGTCCTTACGAGGGGATCGCCGT
GTTCGATGGAAAGAAGATCACGGTCACGGGGACGTTATGGAATGGTAACAAAATCATCGATGAGCGTTTAATTACGC
CGGACGGCTCGATGTTATTCCGTGTTACAATCAATAGCTGA

Corresponding to:

MAHHHHHHENLYFQ↓GSVTGYRLFEEILGSDKEWILQKIYEIMRLLDELGHAEASMRVSDLIYEFMKKGDERLLEEAERLLE
EVERMVFTLEDFVGDWEQTAAYNLDQVLEQGGVSSLLQNLAVSVTPIQRIVRSGENALKIDIHVIIPYEGLSADQMAQIEEV
FKVVYPVDDHHFKVILPYGTLVIDGVTPNMLNYFGRPYEGIAVFDGKKITVTGTLWNGNKIIDERLITPDGSMLFRVTINS-
(Green: 6xHis, Purple: TEVcs, Red: SmBiT, Orange: LCB1, Blue: LgBiT)

**<u>6xHis-TEVcs-SmBiT-1aa-LCB1-0aa-LgBiT (S-BAT)</u>**

ATGGCGCATCACCACCACCACCACGAGAATCTTTATTTCCAAGGTTCGGTGACCGGGTACCGTTTGTTCGAGGAGAT
TCTGGGTGATAAGGAATGGATTTTGCAAAAGATCTATGAGATTATGCGTTTACTTGACGAACTTGGACACGCTGAGGC
GTCGATGCGTGTGTCGGACCTCATTTATGAGTTCATGAAGAAGGGCGACGAGCGCTTGTTGGAAGAGGCCGAGCGT
CTCCTCGAAGAGGTCGAGCGCATGGTATTTACGTTGGAAGATTTCGTGGGAGACTGGGAGCAAACCGCGGCCTATA
ACTTAGATCAGGTGTTGGAGCAAGGTGGTGTTTCGAGTTTACTGCAGAACCTTGCCGTGTCCGTCACACCTATTCAG
CGTATCGTCCGTTCAGGTGAGAATGCACTCAAGATTGACATTCACGTCATCATCCCGTACGAGGGGTTAAGCGCAGA
CCAAATGGCTCAAATCGAAGAGGTATTTAAGGTCGTGTATCCAGTCGATGACCACCATTTTAAAGTCATCCTGCCTTA
TGGCACATTGGTGATTGACGGGGTTACACCAAATATGTTGAACTATTTTGGACGTCCTTACGAGGGGATCGCCGTGTT
CGATGGAAAGAAGATCACGGTCACGGGGACGTTATGGAATGGTAACAAAATCATCGATGAGCGTTTAATTACGCCGG
ACGGCTCGATGTTATTCCGTGTTACAATCAATAGCTGA

Corresponding to:

MAHHHHHHENLYFQ↓GSVTGYRLFEEILGDKEWILQKIYEIMRLLDELGHAEASMRVSDLIYEFMKKGDERLLEEAERLLEE
VERMVFTLEDFVGDWEQTAAYNLDQVLEQGGVSSLLQNLAVSVTPIQRIVRSGENALKIDIHVIIPYEGLSADQMAQIEEVF
KVVYPVDDHHFKVILPYGTLVIDGVTPNMLNYFGRPYEGIAVFDGKKITVTGTLWNGNKIIDERLITPDGSMLFRVTINS-
(Green: 6xHis, Purple: TEVcs, Red: SmBiT, Orange: LCB1, Blue: LgBiT)

**6xHis-TEVcs-SmBiT-1aa-LCB1*-0aa-LgBiT (S-BAT*)**

ATGGCGCATCACCACCACCACCACGAGAATCTTTATTTCCAAGGTTCGGTGACCGGGTACCGTTTGTTCGAGGAGAT
TCTGGGTGATAAGGAATGGATTTTGCAAAAGATCTATGAGATTATGCGTTTACTTGACGAACTTGGACACGCTGAGGC
GTCGATGCGTGTGTCG**GCG**CTCATTTATGAGTTCATGAAGAAGGGCGACGAGCGCTTGTTGGAAGAGGCCGAGCGT
CTCCTCGAAGAGGTCGAGCGCATGGTATTTACGTTGGAAGATTTCGTGGGAGACTGGGAGCAAACCGCGGCCTATA
ACTTAGATCAGGTGTTGGAGCAAGGTGGTGTTTCGAGTTTACTGCAGAACCTTGCCGTGTCCGTCACACCTATTCAG
CGTATCGTCCGTTCAGGTGAGAATGCACTCAAGATTGACATTCACGTCATCATCCCGTACGAGGGGTTAAGCGCAGA
CCAAATGGCTCAAATCGAAGAGGTATTTAAGGTCGTGTATCCAGTCGATGACCACCATTTTAAAGTCATCCTGCCTTA
TGGCACATTGGTGATTGACGGGGTTACACCAAATATGTTGAACTATTTTGGACGTCCTTACGAGGGGATCGCCGTGTT
CGATGGAAAGAAGATCACGGTCACGGGGACGTTATGGAATGGTAACAAAATCATCGATGAGCGTTTAATTACGCCGG
ACGGCTCGATGTTATTCCGTGTTACAATCAATAGCTGA

Corresponding to:

MAHHHHHHENLYFQ↓GSVTGYRLFEEILGDKEWILQKIYEIMRLLDELGHAEASMRVS**A**LIYEFMKKGDERLLEEAERLLEE
VERMVFTLEDFVGDWEQTAAYNLDQVLEQGGVSSLLQNLAVSVTPIQRIVRSGENALKIDIHVIIPYEGLSADQMAQIEEVF
KVVYPVDDHHFKVILPYGTLVIDGVTPNMLNYFGRPYEGIAVFDGKKITVTGTLWNGNKIIDERLITPDGSMLFRVTINS-
(Green: 6xHis, Purple: TEVcs, Red: SmBiT, Orange: LCB1*, Blue: LgBiT)

**6xHis-TEVcs-SmBiT-1aa-LCB1-0aa-LgBiT (S16C)**

ATGGCGCATCACCACCACCACCACGAGAATCTTTATTTCCAAGGT**TGC**GTGACCGGGTACCGTTTGTTCGAGGAGAT
TCTGGGTGATAAGGAATGGATTTTGCAAAAGATCTATGAGATTATGCGTTTACTTGACGAACTTGGACACGCTGAGGC
GTCGATGCGTGTGTCGGACCTCATTTATGAGTTCATGAAGAAGGGCGACGAGCGCTTGTTGGAAGAGGCCGAGCGT
CTCCTCGAAGAGGTCGAGCGCATGGTATTTACGTTGGAAGATTTCGTGGGAGACTGGGAGCAAACCGCGGCCTATA
ACTTAGATCAGGTGTTGGAGCAAGGTGGTGTTTCGAGTTTACTGCAGAACCTTGCCGTGTCCGTCACACCTATTCAG
CGTATCGTCCGTTCAGGTGAGAATGCACTCAAGATTGACATTCACGTCATCATCCCGTACGAGGGGTTAAGCGCAGA
CCAAATGGCTCAAATCGAAGAGGTATTTAAGGTCGTGTATCCAGTCGATGACCACCATTTTAAAGTCATCCTGCCTTA
TGGCACATTGGTGATTGACGGGGTTACACCAAATATGTTGAACTATTTTGGACGTCCTTACGAGGGGATCGCCGTGTT
CGATGGAAAGAAGATCACGGTCACGGGGACGTTATGGAATGGTAACAAAATCATCGATGAGCGTTTAATTACGCCGG
ACGGCTCGATGTTATTCCGTGTTACAATCAATAGCTGA

Corresponding to:

MAHHHHHHENLYFQ↓G**C**VTGYRLFEEILGDKEWILQKIYEIMRLLDELGHAEASMRVSDLIYEFMKKGDERLLEEAERLLEE
VERMVFTLEDFVGDWEQTAAYNLDQVLEQGGVSSLLQNLAVSVTPIQRIVRSGENALKIDIHVIIPYEGLSADQMAQIEEVF
KVVYPVDDHHFKVILPYGTLVIDGVTPNMLNYFGRPYEGIAVFDGKKITVTGTLWNGNKIIDERLITPDGSMLFRVTINS-
(Green: 6xHis, Purple: TEVcs, Red: SmBiT, Orange: LCB1, Blue: LgBiT)

**6xHis-TEVcs-SmBiT-1aa-LCB1-0aa-LgBiT (R55C)**

ATGGCGCATCACCACCACCACCACGAGAATCTTTATTTCCAAGGTTCGGTGACCGGGTACCGTTTGTTCGAGGAGAT
TCTGGGTGATAAGGAATGGATTTTGCAAAAGATCTATGAGATTATGCGTTTACTTGACGAACTTGGACACGCTGAGGC
GTCGATG**TGC**GTGTCGGACCTCATTTATGAGTTCATGAAGAAGGGCGACGAGCGCTTGTTGGAAGAGGCCGAGCGT
CTCCTCGAAGAGGTCGAGCGCATGGTATTTACGTTGGAAGATTTCGTGGGAGACTGGGAGCAAACCGCGGCCTATA
ACTTAGATCAGGTGTTGGAGCAAGGTGGTGTTTCGAGTTTACTGCAGAACCTTGCCGTGTCCGTCACACCTATTCAG
CGTATCGTCCGTTCAGGTGAGAATGCACTCAAGATTGACATTCACGTCATCATCCCGTACGAGGGGTTAAGCGCAGA
CCAAATGGCTCAAATCGAAGAGGTATTTAAGGTCGTGTATCCAGTCGATGACCACCATTTTAAAGTCATCCTGCCTTA
TGGCACATTGGTGATTGACGGGGTTACACCAAATATGTTGAACTATTTTGGACGTCCTTACGAGGGGATCGCCGTGTT
CGATGGAAAGAAGATCACGGTCACGGGGACGTTATGGAATGGTAACAAAATCATCGATGAGCGTTTAATTACGCCGG
ACGGCTCGATGTTATTCCGTGTTACAATCAATAGCTGA

Corresponding to:

MAHHHHHHENLYFQ↓GSVTGYRLFEEILGDKEWILQKIYEIMRLLDELGHAEASM**C**VSDLIYEFMKKGDERLLEEAERLLEE
VERMVFTLEDFVGDWEQTAAYNLDQVLEQGGVSSLLQNLAVSVTPIQRIVRSGENALKIDIHVIIPYEGLSADQMAQIEEVF
KVVYPVDDHHFKVILPYGTLVIDGVTPNMLNYFGRPYEGIAVFDGKKITVTGTLWNGNKIIDERLITPDGSMLFRVTINS-
(Green: 6xHis, Purple: TEVcs, Red: SmBiT, Orange: LCB1, Blue: LgBiT)

**6xHis-TEVcs-SmBiT-1aa-LCB1-0aa-LgBiT (K66C)**

ATGGCGCATCACCACCACCACCACGAGAATCTTTATTTCCAAGGTTCGGTGACCGGGTACCGTTTGTTCGAGGAGAT
TCTGGGTGATAAGGAATGGATTTTGCAAAAGATCTATGAGATTATGCGTTTACTTGACGAACTTGGACACGCTGAGGC
GTCGATGCGTGTGTCGGACCTCATTTATGAGTTCATGAAG**TGC**GGCGACGAGCGCTTGTTGGAAGAGGCCGAGCGT
CTCCTCGAAGAGGTCGAGCGCATGGTATTTACGTTGGAAGATTTCGTGGGAGACTGGGAGCAAACCGCGGCCTATA
ACTTAGATCAGGTGTTGGAGCAAGGTGGTGTTTCGAGTTTACTGCAGAACCTTGCCGTGTCCGTCACACCTATTCAG

CGTATCGTCCGTTCAGGTGAGAATGCACTCAAGATTGACATTCACGTCATCATCCCGTACGAGGGGTTAAGCGCAGA
CCAAATGGCTCAAATCGAAGAGGTATTTAAGGTCGTGTATCCAGTCGATGACCACCATTTTAAAGTCATCCTGCCTTA
TGGCACATTGGTGATTGACGGGGTTACACCAAATATGTTGAACTATTTTGGACGTCCTTACGAGGGGATCGCCGTGTT
CGATGGAAAGAAGATCACGGTCACGGGGACGTTATGGAATGGTAACAAAATCATCGATGAGCGTTTAATTACGCCGG
ACGGCTCGATGTTATTCCGTGTTACAATCAATAGCTGA
Corresponding to:
MAHHHHHHENLYFQ↓GSVTGYRLFEEILGDKEWILQKIYEIMRLLDELGHAEASMRVSDLIYEFMK**C**GDERLLEEAERLLEE
VERMVFTLEDFVGDWEQTAAYNLDQVLEQGGVSSLLQNLAVSVTPIQRIVRSGENALKIDIHVIIPYEGLSADQMAQIEEVF
KVVYPVDDHHFKVILPYGTLVIDGVTPNMLNYFGRPYEGIAVFDGKKITVTGTLWNGNKIIDERLITPDGSMLFRVTINS-
(Green: 6xHis, Purple: TEVcs, Red: SmBiT, Orange: LCB1, Blue: LgBiT)

**6xHis-TEVcs-SmBiT-1aa-LCB1-0aa-LgBiT (R77C)**
ATGGCGCATCACCACCACCACCACGAGAATCTTTATTTCCAAGGTTCGGTGACCGGGTACCGTTTGTTCGAGGAGAT
TCTGGGTGATAAGGAATGGATTTTGCAAAAGATCTATGAGATTATGCGTTTACTTGACGAACTTGGACACGCTGAGGC
GTCGATGCGTGTGTCGGACCTCATTTATGAGTTCATGAAGAAGGGCGACGAGCGCTTGTTGGAAGAGGCCGAG**TGC**
CTCCTCGAAGAGGTCGAGCGCATGGTATTTACGTTGGAAGATTTCGTGGGAGACTGGGAGCAAACCGCGGCCTATA
ACTTAGATCAGGTGTTGGAGCAAGGTGGTGTTTCGAGTTTACTGCAGAACCTTGCCGTGTCCGTCACACCTATTCAG
CGTATCGTCCGTTCAGGTGAGAATGCACTCAAGATTGACATTCACGTCATCATCCCGTACGAGGGGTTAAGCGCAGA
CCAAATGGCTCAAATCGAAGAGGTATTTAAGGTCGTGTATCCAGTCGATGACCACCATTTTAAAGTCATCCTGCCTTA
TGGCACATTGGTGATTGACGGGGTTACACCAAATATGTTGAACTATTTTGGACGTCCTTACGAGGGGATCGCCGTGTT
CGATGGAAAGAAGATCACGGTCACGGGGACGTTATGGAATGGTAACAAAATCATCGATGAGCGTTTAATTACGCCGG
ACGGCTCGATGTTATTCCGTGTTACAATCAATAGCTGA
Corresponding to:
MAHHHHHHENLYFQ↓GSVTGYRLFEEILGDKEWILQKIYEIMRLLDELGHAEASMRVSDLIYEFMKKGDERLLEEAE**C**LLEE
VERMVFTLEDFVGDWEQTAAYNLDQVLEQGGVSSLLQNLAVSVTPIQRIVRSGENALKIDIHVIIPYEGLSADQMAQIEEVF
KVVYPVDDHHFKVILPYGTLVIDGVTPNMLNYFGRPYEGIAVFDGKKITVTGTLWNGNKIIDERLITPDGSMLFRVTINS-
(Green: 6xHis, Purple: TEVcs, Red: SmBiT, Orange: LCB1, Blue: LgBiT)

**6xHis-TEVcs-SmBiT-1aa-LCB1-0aa-LgBiT (S243C)**
ATGGCGCATCACCACCACCACCACGAGAATCTTTATTTCCAAGGTTCGGTGACCGGGTACCGTTTGTTCGAGGAGAT
TCTGGGTGATAAGGAATGGATTTTGCAAAAGATCTATGAGATTATGCGTTTACTTGACGAACTTGGACACGCTGAGGC
GTCGATGCGTGTGTCGGACCTCATTTATGAGTTCATGAAGAAGGGCGACGAGCGCTTGTTGGAAGAGGCCGAGCGT
CTCCTCGAAGAGGTCGAGCGCATGGTATTTACGTTGGAAGATTTCGTGGGAGACTGGGAGCAAACCGCGGCCTATA
ACTTAGATCAGGTGTTGGAGCAAGGTGGTGTTTCGAGTTTACTGCAGAACCTTGCCGTGTCCGTCACACCTATTCAG
CGTATCGTCCGTTCAGGTGAGAATGCACTCAAGATTGACATTCACGTCATCATCCCGTACGAGGGGTTAAGCGCAGA
CCAAATGGCTCAAATCGAAGAGGTATTTAAGGTCGTGTATCCAGTCGATGACCACCATTTTAAAGTCATCCTGCCTTA
TGGCACATTGGTGATTGACGGGGTTACACCAAATATGTTGAACTATTTTGGACGTCCTTACGAGGGGATCGCCGTGTT
CGATGGAAAGAAGATCACGGTCACGGGGACGTTATGGAATGGTAACAAAATCATCGATGAGCGTTTAATTACGCCGG
ACGGCTCGATGTTATTCCGTGTTACAATCAAT**TGT**TGA
Corresponding to:
MAHHHHHHENLYFQ↓GSVTGYRLFEEILGDKEWILQKIYEIMRLLDELGHAEASMRVSDLIYEFMKKGDERLLEEAERLLEE
VERMVFTLEDFVGDWEQTAAYNLDQVLEQGGVSSLLQNLAVSVTPIQRIVRSGENALKIDIHVIIPYEGLSADQMAQIEEVF
KVVYPVDDHHFKVILPYGTLVIDGVTPNMLNYFGRPYEGIAVFDGKKITVTGTLWNGNKIIDERLITPDGSMLFRVTIN**C**-
(Green: 6xHis, Purple: TEVcs, Red: SmBiT, Orange: LCB1, Blue: LgBiT)

**SmBiT-1aa-LCB1-0aa-LgBiT (tag-free S-BAT)**
ATGGGTTCGGTGACCGGGTACCGTTTGTTCGAGGAGATTCTGGGTGATAAGGAATGGATTTTGCAAAAGATCTATGA
GATTATGCGTTTACTTGACGAACTTGGACACGCTGAGGCGTCGATGCGTGTGTCGGACCTCATTTATGAGTTCATGAA
GAAGGGCGACGAGCGCTTGTTGGAAGAGGCCGAGCGTCTCCTCGAAGAGGTCGAGCGCATGGTATTTACGTTGGAA
GATTTCGTGGGAGACTGGGAGCAAACCGCGGCCTATAACTTAGATCAGGTGTTGGAGCAAGGTGGTGTTTCGAGTTT
ACTGCAGAACCTTGCCGTGTCCGTCACACCTATTCAGCGTATCGTCCGTTCAGGTGAGAATGCACTCAAGATTGACA
TTCACGTCATCATCCCGTACGAGGGGTTAAGCGCAGACCAAATGGCTCAAATCGAAGAGGTATTTAAGGTCGTGTAT
CCAGTCGATGACCACCATTTTAAAGTCATCCTGCCTTATGGCACATTGGTGATTGACGGGGTTACACCAAATATGTTG
AACTATTTTGGACGTCCTTACGAGGGGATCGCCGTGTTCGATGGAAAGAAGATCACGGTCACGGGGACGTTATGGAA
TGGTAACAAAATCATCGATGAGCGTTTAATTACGCCGGACGGCTCGATGTTATTCCGTGTTACAATCAATAGCTGA
Corresponding to:
MGSVTGYRLFEEILGDKEWILQKIYEIMRLLDELGHAEASMRVSDLIYEFMKKGDERLLEEAERLLEEVERMVFTLEDFVGD
WEQTAAYNLDQVLEQGGVSSLLQNLAVSVTPIQRIVRSGENALKIDIHVIIPYEGLSADQMAQIEEVFKVVYPVDDHHFKVIL
PYGTLVIDGVTPNMLNYFGRPYEGIAVFDGKKITVTGTLWNGNKIIDERLITPDGSMLFRVTINS-
(Red: SmBiT, Orange: LCB1, Blue: LgBiT)

**SmBiT-1aa-LCB1*-0aa-LgBiT (tag-free S-BAT*)**

ATGGGTTCGGTGACCGGGTACCGTTTGTTCGAGGAGATTCTGGGTGATAAGGAATGGATTTTGCAAAAGATCTATGA
GATTATGCGTTTACTTGACGAACTTGGACACGCTGAGGCGTCGATGCGTGTGTCG**GCG**CTCATTTATGAGTTCATGA
AGAAGGGCGACGAGCGCTTGTTGGAAGAGGCCGAGCGTCTCCTCGAAGAGGTCGAGCGCATGGTATTTACGTTGGA
AGATTTCGTGGGAGACTGGGAGCAAACCGCGGCCTATAACTTAGATCAGGTGTTGGAGCAAGGTGGTGTTTCGAGTT
TACTGCAGAACCTTGCCGTGTCCGTCACACCTATTCAGCGTATCGTCCGTTCAGGTGAGAATGCACTCAAGATTGAC
ATTCACGTCATCATCCCGTACGAGGGGTTAAGCGCAGACCAAATGGCTCAAATCGAAGAGGTATTTAAGGTCGTGTA
TCCAGTCGATGACCACCATTTTAAAGTCATCCTGCCTTATGGCACATTGGTGATTGACGGGGTTACACCAAATATGTT
GAACTATTTTGGACGTCCTTACGAGGGGATCGCCGTGTTCGATGGAAAGAAGATCACGGTCACGGGGACGTTATGG
AATGGTAACAAAATCATCGATGAGCGTTTAATTACGCCGGACGGCTCGATGTTATTCCGTGTTACAATCAATAGCTGA
Corresponding to:
MGSVTGYRLFEEILGDKEWILQKIYEIMRLLDELGHAEASMRVS**A**LIYEFMKKGDERLLEEAERLLEEVERMVFTLEDFVGD
WEQTAAYNLDQVLEQGGVSSLLQNLAVSVTPIQRIVRSGENALKIDIHVIIPYEGLSADQMAQIEEVFKVVYPVDDHHFKVIL
PYGTLVIDGVTPNMLNYFGRPYEGIAVFDGKKITVTGTLWNGNKIIDERLITPDGSMLFRVTINS-
(Red: SmBiT, Orange: LCB1*, Blue: LgBiT)

**Aga2p Signal Sequence-SmBiT-1aa-LCB1-0aa-LgBiT (yeast secretion S-BAT)**

atgcagttacttcgctgtttttcaatattttctgttattgcttcagttttagcaGGTTCGGTGACCGGGTACCGTTTGTTCGAGGAGATTCTGGGTGAT
AAGGAATGGATTTTGCAAAAGATCTATGAGATTATGCGTTTACTTGACGAACTTGGACACGCTGAGGCGTCGATGCGT
GTGTCGGACCTCATTTATGAGTTCATGAAGAAGGGCGACGAGCGCTTGTTGGAAGAGGCCGAGCGTCTCCTCGAAG
AGGTCGAGCGCATGGTATTTACGTTGGAAGATTTCGTGGGAGACTGGGAGCAAACCGCGGCCTATAACTTAGATCAG
GTGTTGGAGCAAGGTGGTGTTTCGAGTTTACTGCAGAACCTTGCCGTGTCCGTCACACCTATTCAGCGTATCGTCCG
TTCAGGTGAGAATGCACTCAAGATTGACATTCACGTCATCATCCCGTACGAGGGGTTAAGCGCAGACCAAATGGCTC
AAATCGAAGAGGTATTTAAGGTCGTGTATCCAGTCGATGACCACCATTTTAAAGTCATCCTGCCTTATGGCACATTGG
TGATTGACGGGGTTACACCAAATATGTTGAACTATTTTGGACGTCCTTACGAGGGGATCGCCGTGTTCGATGGAAAG
AAGATCACGGTCACGGGGACGTTATGGAATGGTAACAAAATCATCGATGAGCGTTTAATTACGCCGGACGGCTCGAT
GTTATTCCGTGTTACAATCAATAGCTGA
Corresponding to:
MQLLRCFSIFSVIASVLAGSVTGYRLFEEILGDKEWILQKIYEIMRLLDELGHAEASMRVSDLIYEFMKKGDERLLEEAERLLE
EVERMVFTLEDFVGDWEQTAAYNLDQVLEQGGVSSLLQNLAVSVTPIQRIVRSGENALKIDIHVIIPYEGLSADQMAQIEEV
FKVVYPVDDHHFKVILPYGTLVIDGVTPNMLNYFGRPYEGIAVFDGKKITVTGTLWNGNKIIDERLITPDGSMLFRVTINS-
(Green: Aga2p Signal Sequence, Red: SmBiT, Orange: LCB1, Blue: LgBiT)

**Aga2p Signal Sequence-SmBiT-1aa-LCB1*-0aa-LgBiT (yeast secretion S-BAT*)**

atgcagttacttcgctgtttttcaatattttctgttattgcttcagttttagcaGGTTCGGTGACCGGGTACCGTTTGTTCGAGGAGATTCTGGGTGAT
AAGGAATGGATTTTGCAAAAGATCTATGAGATTATGCGTTTACTTGACGAACTTGGACACGCTGAGGCGTCGATGCGT
GTGTCG**GCG**CTCATTTATGAGTTCATGAAGAAGGGCGACGAGCGCTTGTTGGAAGAGGCCGAGCGTCTCCTCGAAG
AGGTCGAGCGCATGGTATTTACGTTGGAAGATTTCGTGGGAGACTGGGAGCAAACCGCGGCCTATAACTTAGATCAG
GTGTTGGAGCAAGGTGGTGTTTCGAGTTTACTGCAGAACCTTGCCGTGTCCGTCACACCTATTCAGCGTATCGTCCG
TTCAGGTGAGAATGCACTCAAGATTGACATTCACGTCATCATCCCGTACGAGGGGTTAAGCGCAGACCAAATGGCTC
AAATCGAAGAGGTATTTAAGGTCGTGTATCCAGTCGATGACCACCATTTTAAAGTCATCCTGCCTTATGGCACATTGG
TGATTGACGGGGTTACACCAAATATGTTGAACTATTTTGGACGTCCTTACGAGGGGATCGCCGTGTTCGATGGAAAG
AAGATCACGGTCACGGGGACGTTATGGAATGGTAACAAAATCATCGATGAGCGTTTAATTACGCCGGACGGCTCGAT
GTTATTCCGTGTTACAATCAATAGCTGA
Corresponding to:
MQLLRCFSIFSVIASVLAGSVTGYRLFEEILGDKEWILQKIYEIMRLLDELGHAEASMRVS**A**LIYEFMKKGDERLLEEAERLLE
EVERMVFTLEDFVGDWEQTAAYNLDQVLEQGGVSSLLQNLAVSVTPIQRIVRSGENALKIDIHVIIPYEGLSADQMAQIEEV
FKVVYPVDDHHFKVILPYGTLVIDGVTPNMLNYFGRPYEGIAVFDGKKITVTGTLWNGNKIIDERLITPDGSMLFRVTINS-
(Green: Aga2p Signal Sequence, Red: SmBiT, Orange: LCB1, Blue: LgBiT)
